# Neurotrophic factor MANF regulates autophagy and lysosome function to promote proteostasis in *C. elegans*

**DOI:** 10.1101/2023.07.31.551399

**Authors:** Shane K. B. Taylor, Jessica H. Hartman, Bhagwati P. Gupta

## Abstract

The conserved mesencephalic astrocyte-derived neurotrophic factor (MANF) protects dopaminergic neurons but also functions in several other tissues. Previously, we showed that *Caenorhabditis elegans manf-1* null mutants have increased ER stress, dopaminergic neurodegeneration, protein aggregation, slower growth, and a reduced lifespan. The multiple requirements of MANF in different systems suggest its essential role in regulating cellular processes. However, how intracellular and extracellular MANF regulates broader cellular function remains unknown. Here, we report a novel mechanism of action for *manf-1* that involves the autophagy transcription factor HLH-30/TFEB-mediated signaling to regulate lysosomal function and aging. We generated multiple transgenic strains overexpressing MANF-1 and found that animals had extended lifespan, reduced protein aggregation, and improved neuronal health. Using a fluorescently tagged MANF-1, we observed different tissue localization of MANF-1 depending on the ER retention signal. Further subcellular analysis showed that MANF-1 localizes within cells to the lysosomes. These findings were consistent with our transcriptomic studies and, together with analysis of autophagy regulators, demonstrate that MANF-1 regulates protein homeostasis through increased autophagy and lysosomal activity. Collectively, our findings establish MANF as a critical regulator of the stress response, proteostasis, and aging.

## INTRODUCTION

Nothing is perfect, and this includes the intracellular machinery of a cell. Despite their constant working, the ability of cells to maintain homeostasis declines with age [1]. Dysregulated homeostasis can cause protein misfolding and the formation of protein clumps or aggregates. Abnormal accumulation of protein aggregates is toxic to cells because it increases cellular stress, disrupts physiological processes, and eventually results in the failure of cells, tissues, and organs to maintain their function. Current research suggests that maintaining protein homeostasis (proteostasis) is essential for functioning cellular processes, reducing age-related diseases, and promoting healthy aging. Notably, cells can activate stress response signaling pathways, such as the unfolded protein response (UPR), to maintain proteostasis and minimize cellular damage. Although these responses work very well in most cases, they are not always efficient. Failure to eliminate toxic protein aggregates could lead to age-related diseases, including neurodegenerative diseases such as Parkinson’s Disease (PD) and Huntington’s Disease (HD) [1–3].

A promising approach to enhance proteostasis and protect against neurodegeneration involves the therapeutic delivery of neurotrophic factors (NTFs). NTFs are proteins that support neuronal survival, growth, and differentiation. Their anti-inflammatory and anti-apoptotic properties make them promising therapeutic candidates [4]. One family of NTFs relevant to this study is represented by two proteins, namely the cerebral dopamine neurotrophic factor (CDNF) and the <u>m</u>esencephalic <u>a</u>strocyte-derived <u>n</u>eurotrophic <u>f</u>actor (MANF) [5]. These two proteins are structurally and mechanistically distinct from other classical NTFs [4, 6]. Although vertebrates express both CDNF and MANF, only MANF is expressed in invertebrates.

MANF homologs have been studied in different animals including mouse, rat, zebrafish, and two leading invertebrates, *Drosophila melanogaster* and *Caenorhabditis elegans*. These studies have shown that MANF confers cytoprotection through endoplasmic reticulum (ER)-UPR regulation [5]. ER-UPR is a quality control mechanism that rectifies cellular stress by correctly folding misfolded proteins, reducing translation, and activating apoptosis when homeostasis cannot be restored [7]. Three conserved transmembrane proteins mediate the ER-UPR, inositol-requiring enzyme 1 (IRE1), protein kinase RNA (PKR)-like ER kinase (PERK), and activating transcription factor 6 (ATF6) (*ire-1*, *pek-1*, and *atf-6*, respectively, in *C. elegans*), which activate overlapping but distinct pathways in stress responses [7, 8]. In addition, IRE1 creates a spliced form of the transcription factor X-box binding protein-1 (XBP1), the main regulator of ER-UPR. Spliced XBP-1 maintains homeostasis through the transcription of chaperones, such as GRP78/BiP (*C. elegans hsp-4*), which is affected by MANF [7–11]. Consistent with the role of MANF in ER-UPR maintenance, its sequence contain an ER retention signal [12]. Under normal conditions, MANF is retained in the ER and is secreted in response to protein misfolding or cellular stress [13].

Recent studies have indicated that MANF is also linked to other diseases with dysregulated homeostasis, more specifically, metabolic dysfunction, diabetes, ischemia, and retinal degeneration [4, 14, 15]. Accordingly, MANF is widely expressed [14–17] underscoring the protein’s critical roles in multiple processes. Although MANF is secreted from the ER in response to stress, how it regulates cellular function and how extracellular MANF affects other tissues is currently unknown. Is there a common theme underlying its mechanisms of action in different cell types? The nematode *Caenorhabditis elegans* offers the necessary genetic and molecular tools to investigate these questions and functions of MANF [18–20]. Studies on the *C. elegans* MANF homolog, MANF-1, have shown that it is secreted and endocytosed upon binding to sulfatides [21]. Our group and others have demonstrated the role of MANF-1 in processes that include age-dependent protection of DA neurons [22], regulation of the ER-UPR chaperone HSP-4/BIP/GRP78, and sensitivity to bacterial pathogenesis [23, 24]. The protective role of *manf-1* may be mediated by genes affecting autophagy, ER-UPR, and immunity [23, 24].

This study reports novel findings on *C. elegans* MANF-1 and the mechanism of function. Our data indicate that *manf-1* mutants died prematurely when subjected to ER stress, and MANF-1 expression was upregulated in response to stress-inducing conditions. Further analysis revealed an essential role for this gene in regulating ER-UPR signaling. We generated multiple transgenic strains overexpressing MANF-1 (*hsp::manf-1*, *manf-1p::manf-1*, and *manf-1p::manf-1::mCherry*) and found that the animals had longer lifespans, improved proteostasis, and reduced neurodegeneration. Consistent with these results, the MANF-1::mCherry chimeric protein was widely expressed and secreted. Studies using tissue-specific and subcellular markers revealed that MANF-1 was localized to areas including the intestine, muscles, coelomocytes, and, more specifically, lysosomes. Further characterization revealed differences in expression depending on the presence or absence of the ER retention signal; however, lysosomal localization was present in all the strains. Moreover, the MANF-1 pattern was broadly similar in animals carrying the native ER sequence (KEEL). Interestingly, the beneficial effects of MANF-1 did not depend on the ER sequence. Consistent with these data, transcroptomic profiling of the mutant and overexpressing animals revealed significant changes in the expression of ER-UPR and lysosomal genes. Further experiments conducted to investigate autophagy in MANF-1 mutant and overexpression animals using the LGG-1 and p62/SQST-1 reporters suggest that MANF-1 regulates autophagic flux. We also observed that MANF-1 overexpression caused HLH-30/TFEB, a transcription factor that regulates autophagy and lysosomal biogenesis, to be upregulated and localized to the nucleus. Among other changes, we observed reduced lipid levels in the transgenic animals, consistent with increased autophagy and lysosomal activity. Altogether, our findings establish MANF-1 as more than a neurotrophic factor, but rather as a player regulating ER-UPR, autophagy, and lysosomal function, potentially in a coordinated manner, to maintain proteostasis, neuronal health, and the lifespan of animals.

## Results

### ER stress activates *manf-1* expression and affects viability and lifespan of mutant animals

We previously characterized the role of *C. elegans manf-1* using mutant strains and a *manf-1p::GFP* transgenic line [22]. Next, *manf-1p::GFP* expression was examined following treatment with the ER stress-activating drug tunicamycin. The results showed a significant increase in GFP fluorescence (Fig. 1A). Consistent with this finding, *manf-1* transcription was upregulated in animals subjected to acute (8 hr) and chronic (3 days) tunicamycin treatment (Fig. 1B and C). We also used an endogenously tagged *mKate2::manf-1* strain [23] and observed that tunicamycin caused a >100-fold increase in chimeric protein abundance (Fig. 1D). Thus, *manf-1* expression responds to ER stress and is upregulated to confer protection.

**Figure 1.**
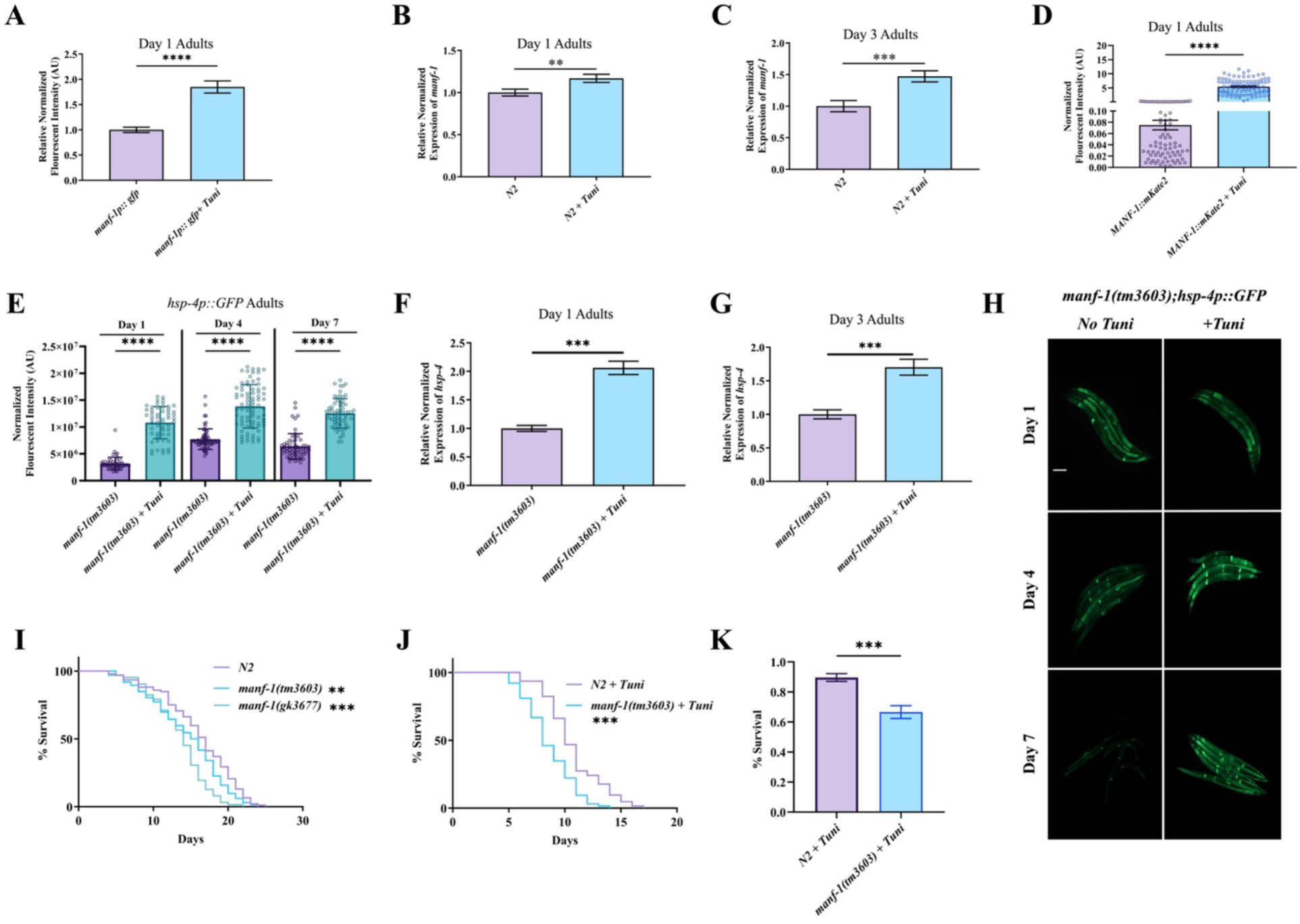
*manf-1* expression and *manf-1(tm3603)* mutant phenotypes in response to increased ER stress. **(A)** Quantification of GFP fluorescence in *manf-1p::GFP* day 1 adults following an 8 hr treatment of 5 μg/mL tunicamycin. **(B)** *manf-*1 RT-qPCR analysis in day 1 N2 adults following an 8 hr treatment of 5 μg/mL tunicamycin. **C)** Same as B, except that animals were chronically exposed to 5 μg/mL tunicamycin until day 3 of adulthood. **D)** Quantification of mKate2 fluorescence in the *manf-1p::manf-1::mKate2* transgenic strain. Day 1 adults were treated with 25ng/μL tunicamycin overnight. **E)** Temporal analysis of GFP fluorescence in *manf-1(tm3603)*; *hsp-4p::GFP* strain with and without 25 ng/µL tunicamycin exposure for 4 hrs on days 1, 4 and 7 of adulthood. **F)** RT-qPCR analysis of *hsp-4* in day 1 *manf-1(tm3603)* adults following 8 hr exposure to 5 μg/mL tunicamycin. **G)** Same as F, except that animals were chronically exposed to 5 µg/mL tunicamycin until day 3 of adulthood. **H)** Representative images of *hsp-4p::GFP* fluorescing animals corresponding to the panel E (scale bar 100 µm). **I, J)** Lifespan analysis of *manf-1* mutants and N2 control without any treatment (I) and following chronic exposure to 25 ng/µL tunicamycin (J). **K)** Percentage survival following 4 hrs exposure to 50 ng/µL tunicamycin in liquid. For results presented in panels A, D, E, I-K, a total of 60 – 140 worms from three independent batches (20 worms minimum per batch) were examined. The RT-qPCR experiments (B, C, F, G) were carried out in three independent batches. The graphs are plotted as mean ± SEM (A-D, F, G, I) and mean ± SD (E). Data was analyzed using student’s t-test (A-I) and the log-rank (Kaplan-Meier) method (J and K). \**p*<0.05; \*\**p*<0.01; \*\*\**p*<0.001; \*\*\*\**p*<0.0001.

The *manf-1* mutants exhibit increased *hsp-4* expression [21–23]. As ER-UPR is known to decline with age [1, 25], we examined the effect of MANF-1 in older *manf-1 (tm3603)* worms by treating them with tunicamycin. Levels of *hsp-4p::GFP* were quantified in 1-, 4-, and 7-day old adults. The results showed that additional ER stress induction in mutant animals using tunicamycin caused further upregulation of both GFP fluorescence and *hsp-4* transcripts with age (Fig. 1E–H). Together with the findings of a previous study that reported higher *hsp-4p::GFP* expression in *manf-1* mutant larvae and day 1 adults [22, 23], these data led us to conclude that *manf-1* is essential for ER-UPR maintenance throughout the lifespan of animals. The crucial role of the protein was demonstrated through additional experiments that showed that mutations in *manf-1* significantly reduced the lifespan of animals (Fig. 1I). Moreover, chronic tunicamycin exposure enhanced DA neurodegeneration (Supplementary Figure 1A) and caused a significant reduction in lifespan (Fig. 1J). Similarly, acute tunicamycin treatment significantly increased the mortality of *manf-1(tm3603)* animals (Fig. 1K).

Finally, we examined whether *manf-1* affected the sensitivity of animals to other forms of stress. Therefore, heat stress and oxidative stress assays were performed in this study. Except for the paraquat, where chronic exposure caused significant damage to DA neurons in *manf-1* mutants (Supplementary Figure 1D), no other difference was observed between mutant and wild-type animals (Supplementary Figure 1B, C). Taken together, the results presented in this section demonstrate that *manf-1* is necessary for the maintenance of ER-UPR and protects animals against ER stress.

### *manf-1* interacts with ER-UPR components and mutants show enhanced protein aggregation defects

Based on the upregulation of *hsp-4* in *manf-1(tm3603)* animals, we investigated the ER-UPR genes that define each of the signaling arms, namely ATF6/*atf-6*, PERK/*pek-1*, IRE1/*ire-1*, and the downstream transcription factor XBP-1/*xbp-1* [7, 11]. The expression of all four genes was significantly upregulated in *manf-1* mutants compared to wild-type controls (Fig. 2A). The ratio of spliced to total *xbp-1* was also high (Fig. 2B). These data suggest that *manf-1* affects a wide range of processes that maintain ER homeostasis.

**Figure 2.**
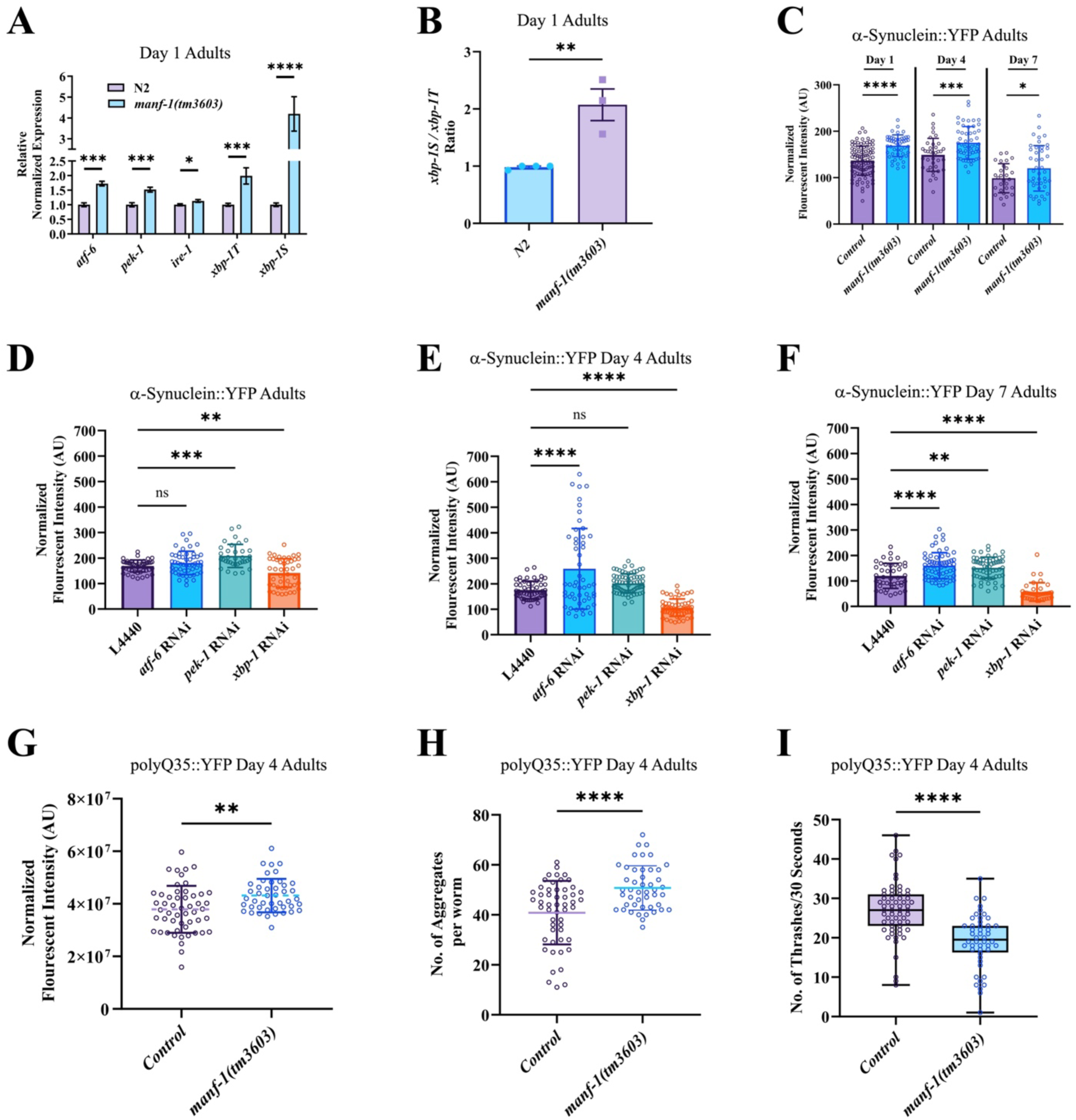
Effect of RNAi knockdown of ER-UPR components on α-Synuclein and polyQ aggregation in *manf-1(tm3603)* animals. **A)** RT-qPCR analysis of ER-UPR genes (*atf-6, pek-1,ire-1* and *xbp-1)* in N2 and *manf-1(tm3603)* day 1 adults. *Xbp-1* total and spliced forms are indicated by *xbp-1T* and *xbp-1S*, respectively. **B)** The ratio of spliced *xbp-1* to total *xbp-1* in *manf-1(tm3603)* day 1 adults. **C)** The fluorescence intensity of α-Synuclein::YFP in body wall muscles was measured in day 1, 4 and 7 adults of *manf-1(tm3603)* and control genotypes. **D-F)** α-Synuclein::YFP fluorescence intensities (D), day 4 (E) and day 7 (F), following RNAi knockdown of *atf-6, pek-1* and *xbp-1*. L4440 is an empty vector RNAi control. **G-I)** polyQ35 protein aggregation phenotype in *manf-1(tm3603)* animals on day 4 of adulthood compared to wild-type. The total fluorescent intensity measurement of polyQ35::YFP (G) and number of aggregates (H) in body wall muscles. The thrashing response of animals over a 30 second interval (I). Results in panels A and B are based on pooled worms from 3 batches. For panels C-J, at least 3 batches were analyzed (10-20 worms per batch). Results are shown as mean ± SEM (A, B) and mean ± SD (C-I). Panel J shows a box plot that contains all data points along with the mean and lower and upper boundaries marking the 25^th^ and 75^th^ quartile, respectively. Data was analyzed using student’s *t*-test (A-C, G-I) and one-way ANOVA with Dunnett’s test (D-F). *p<0.05; **p<0.01; ***p<0.001; ****p<0.0001. ns: not significant.

As ER stress affects the ability of cells to promote protein folding and reduce the accumulation of misfolded and unfolded proteins, we examined the proteostasis defects in *manf-1* mutants. To this end, transgenic animals expressing YFP reporter-tagged human α-Synuclein in body wall muscles were used. In a previous study, we demonstrated that these transgenic animals showed increased aggregation in the absence of *manf-1* [22]. Analysis of α-Synuclein::YFP in 1-, 4-, and 7-day old *manf-1 (tm3603)* adults revealed higher aggregation compared with that in wild-type controls (Fig. 2C, Supplementary Figure 2A), suggesting that *manf-1* is necessary to maintain proteostasis at different stages of adulthood. The phenotype was further enhanced by RNAi knockdown of *atf-6* and *pek-1* (Fig. 2D–F and Supplementary Figure 2A), providing evidence that ER-UPR was compromised but not eliminated in the absence of *manf-1*. Notably, the *manf-1(tm3603); xbp-1*(RNAi) animals showed an opposite effect, i.e., α-Synuclein::YFP levels were considerably reduced (Fig. 2D–F and Supplementary Figure 2A). Although the precise reasons are unknown, we noted that the animals were unhealthy and exhibited slower growth, abnormal movement, reduced viability, and a tendency to frequently burst open through the vulval opening (see Methods) (Supplementary Figure 3). These observations suggest that *xbp-1* knockdown significantly compromised the physiological processes in *manf-1* mutants, leading to reduced α-Synuclein aggregation. The synthetic interaction between *manf-1* and *xbp-1* was consistent with that observed in a previous study that reported that *manf-1; ire-1* double mutants and *xbp-1* RNAi-treated *manf-1* mutants were lethal or sterile [23].

We used another protein aggregation system to examine the effects of *manf-1*, which consists of glutamine repeats (polyQ) and serves as a *C. elegans* HD model [26]. To this end, two polyQ strains, Q35::YFP (AM140) and Q40::YFP (AM141), that express fusion proteins in body wall muscles under the *unc-54* promoter were utilized [26]. Similar to α-Synuclein, both polyQ carrying worms exhibited a significant increase in protein aggregation in *manf-1* mutants (Fig. 2G, H and Supplementary Figure 2B, C). Thrashing defects in the polyQ35 animals, due to aggregates accumulating in muscles, were also enhanced (Fig. 2I). Thus, *manf-1* is necessary for regulating ER-UPR signaling and preventing protein aggregation.

### Overexpression of *manf-1* reduces protein aggregation, improves neuronal survival, and extends lifespan

The essential role of *manf-1* in stress response maintenance and proper protein folding raises the possibility that overexpression of this gene in animals may have beneficial and protective effects. To test this hypothesis, we used different approaches to activate *manf-1* expression. One of these involves treating animals with bioactive compounds such as curcumin and lithium. Recently, these two chemicals were reported to increase *manf* transcription in mammalian cells [27, 28]. Similar experiments in worms also showed increased *manf-1* levels, albeit only modestly (SKBT, unpublished). Other approaches involve the use of transgenic strains that express *manf-1* under the control of either the heat-shock *hsp-16.41* promoter *(hsp::manf-1)* or the native promoter (*manf-1p::manf-1* and *manf-1p::manf-1::mCherry*) (see Methods).

For the *hsp::manf-1* strain, we tested whether induction *manf-1* expression had a cytoprotective effect in animals. The transgenic worms showed a robust response to heat treatments because a 1 hr 31-°C exposure caused a significant increase in *manf-1* transcription (Fig. 3A). Notably, heat shock was not required at later stages as 3- and 7-day old adults showed a significant expression without any treatment (Fig. 3B), likely due to basal activity of the promoter at room temperature.

**Figure 3.**
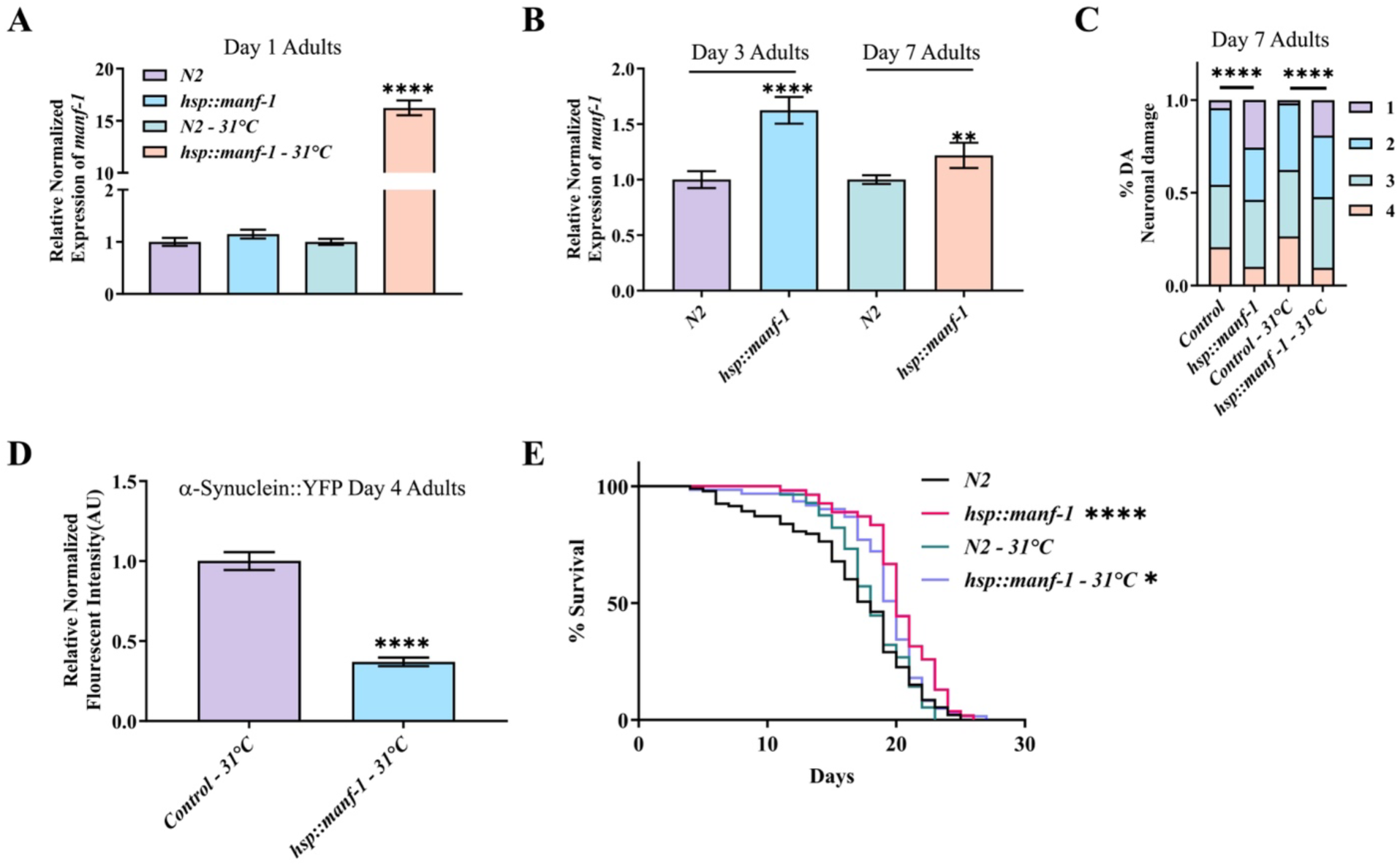
Effect of *manf-1* overexpression on animals using a *hsp::manf-1* system. **A)** RT-qPCR analysis of *manf-1* expression in day 1 *hsp::manf-1* and N2 animals. Animals were maintained either at 20°C or subjected to 1 hr heat shock at 31°C. **B)** RT-qPCR analysis of day 3 and day 7 *hsp::manf-1* and N2 adults grown at 20°C. **C)** Neuronal analysis in day 7 *hsp::manf-1* adults. An odd day heat treatment at 31°C starting on day 1 until day 7 was used to activate *hsp::manf-*Neurons were scored as described in Methods (1: normal cell bodies and dendrites, 2: dendritic damage, 3: cell body missing or abnormally shaped, 4: both dendrites and cell bodies defective). **D)** The effect of *hsp::manf-1* overexpression on α-Synuclein aggregation. The fluorescent intensity of α-Synuclein::YFP was measured in day 7 *hsp::manf-1* adults following the same heat treatment as in panel C. **E)** The lifespan analysis of *hsp::manf-1* animals after being subjected to the same heat treatment as in panels C and D. Data shown in panels A-C include three different batches. For α-Synuclein and lifespan analyses, three batches with 20-30 animals per batch were examined. Results are expressed as mean ± SEM (A, B, D). Data was analyzed using one-way ANOVA with Tukey’s test (A, B). Chi square test (C), Student’s *t*-test (D), and log-rank (Kaplan-Meier) method (E). \**p*<0.05; \*\**p*<0.01; \*\*\**p*<0.001; \*\*\*\**p*<0.0001.

Phenotypic analysis of day 7 *hsp::manf-1; dat-1p::YFP* adults showed increased protection of DA neurons following heat treatment. Specifically, these animals had a much higher proportion of morphologically normal dendritic processes and cell bodies (Fig. 3C). Quantification of neuronal defects revealed fewer animals with defective dendritic and cell body morphologies, demonstrating the beneficial effects of *manf-1* in promoting neuronal health. Interestingly, animals without heat treatment also showed comparable neuroprotection (Fig. 3C), suggesting that a mild increase in MANF-1 levels is sufficient to protect DA neurons. In addition, the *hsp::manf-1* transgene caused a significantly lower α-Synuclein::YFP fluorescence in *hsp::manf-1; α-Synuclein::YFP* animals, suggesting a reduction in protein aggregation defects in older adults (Fig. 3D). Finally, the lifespan was significantly enhanced (approximately 5% in heat-treated animals and 21% in untreated animals; Fig. 3E).

Next, we examined the phenotype of *manf-1p::MANF-1::mCherry* transgenic animals (*MANF-1^KEEL::mCherry^*). The *MANF-1^KEEL::mCherry^* worms showed high levels of secreted MANF-1 in three different lines examined (see next section and Methods). One of these lines showing high transgene transmission (DY759, *bhEx304*) was analyzed in detail. While *hsp-4* transcripts were not significantly affected in these animals, the *hsp-4p::GFP* transgene was activated resulting in increased GFP fluorescence, which suggests that *manf-1* overexpression caused a modest activation of ER-UPR (Fig. 4A and B). We also observed an increase in both the total and spliced *xbp-1* transcripts, whereas *pek-1* levels were reduced (Supplementary Figure 4). However, *MANF-1^KEEL::mCherry^* suppressed the ER stress phenotype of *manf-1(tm3603)* mutants based on *hsp-4::GFP* analysis (Fig. 4A and B).

**Figure 4.**
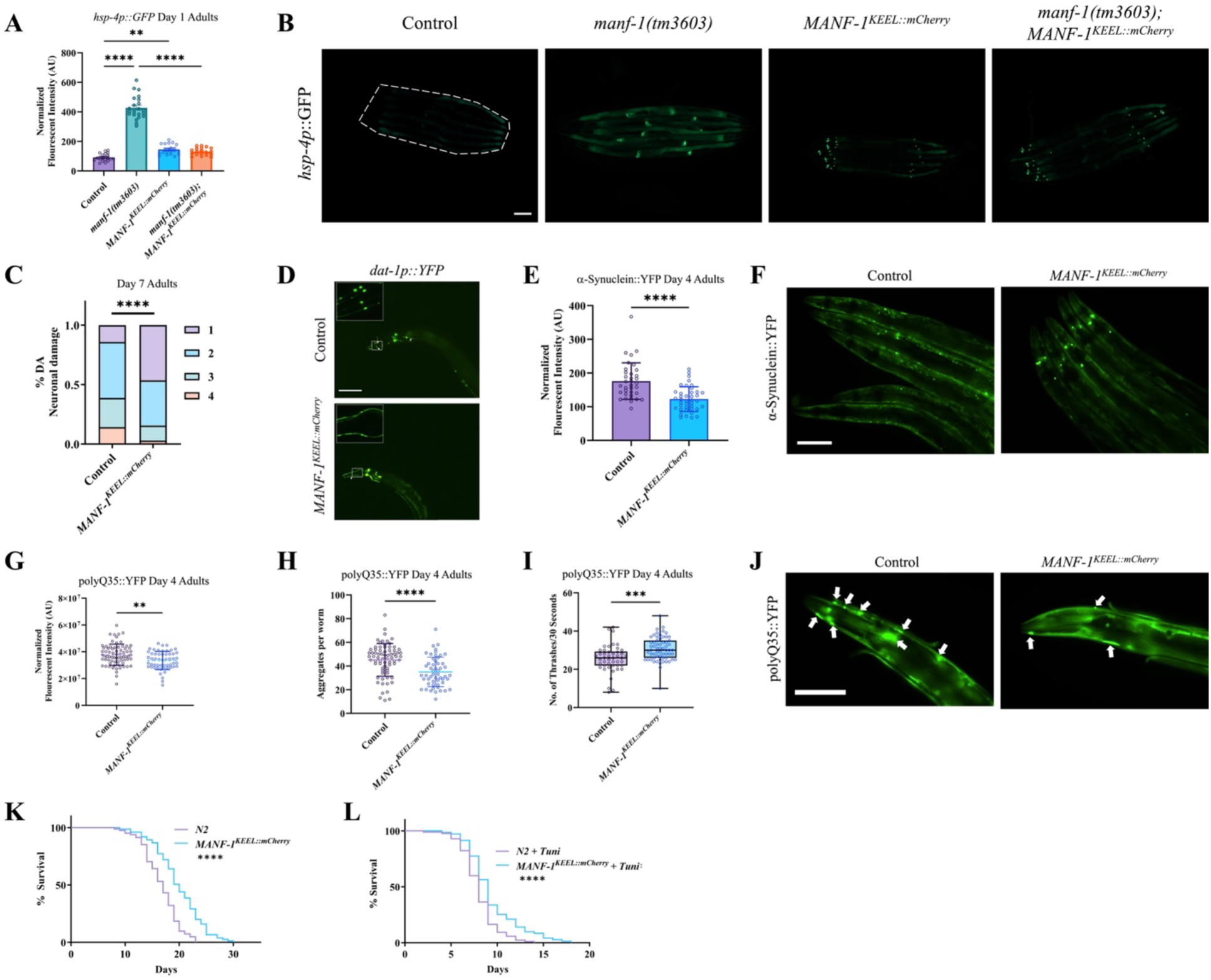
Effect of *manf-1* overexpression on neurons, stress response, and lifespan of animals. **A-B)** *hsp-4p::GFP* reporter expression in wild-type, *manf-1(tm3603)*, *MANF-1^KEEL::mCherry^*and *manf-1(tm3603); MANF-1^KEEL::mCherry^* day 1 adults. Quantification of *hsp-4p::*GFP fluorescence (A) and representative images (scale bar 100 µm) (B). **C, D)** Analysis of dopaminergic neurons in day 7 *MANF-1^KEEL::mCherry^*adults. (C) Neurons were classified into four different categories based on defects in cell bodies and dendrites. See Figure 3 legend and Methods for details. (D) Corresponding images of dopaminergic neurons (scale bar 100 µm). **E, F)** The effect of *MANF-1^KEEL::mCherry^*on α-Synuclein::YFP aggregation in body wall muscles of day 4 adults. (E) The α-Synuclein::YFP fluorescence intensity was measured in day 4 *MANF-1^KEEL::mCherry^*adults; (F) Representative images of α-Synuclein::YFP fluorescence in control and *MANF-1^KEEL::mCherry^* animals at day 4 of adulthood (scale bar 100 µm). **G-J)** The effect of *MANF-1^KEEL::mCherry^*on polyQ35::YFP aggregation in body wall muscles of day 4 adults. Quantification of polyQ35::YFP fluorescence intensity (G) and polyQ35::YFP protein aggregates (H). (I) Thrashing response of animals over a 30 sec interval. (J) Representative images of day 4 polyQ35::YFP adult animals (scale bar 100 µm). Arrows mark aggregates in the head region of worm. **K, L)** Lifespan analysis of wild-type N2 and *MANF-1^KEEL::mCherry^* transgenic animals without any treatment (K) and in the presence of 25 ng/μL tunicamycin (L). At least three batches with 10-30 worms per batch were examined for panels A-I and three batches with 20-30 animals per batch for panels K, L. Data in A is expressed as mean ± SEM and in E-H as mean ± SD. Panel I shows a box plot containing all data points along with the mean and lower and upper boundaries marking the 25^th^ and 75^th^ quartile, respectively. Data was analyzed using one-way ANOVA with Tukey’s test (A), Chi-square test (C), Student’s *t*-test (E-J), and log-rank (Kaplan-Meier) method (K & L). *p<0.05; **p<0.01; ***p<0.001; ****p<0.0001.

To determine if *MANF-1^KEEL::mCherry^* had protective capabilities, we analyzed the DA neuron morphology, α-Synuclein and polyQ aggregation, movement, and lifespan of animals. The results indicated that *manf-1* overexpression conferred significant protection on DA neurons (Fig. 4D) and reduced the aggregation of α-Synuclein (Fig. 4E) and polyQ35 & polyQ40 (Fig. 4G, H, J and Supplementary Figure 2C). Additionally, the thrashing response of muscular polyQ35-expressing worms significantly improved (Fig. 4I). Similar to *hsp::manf-1* animals; the *MANF-1^KEEL::mCherry^* animals had a significantly increased lifespan (19.2% higher; mean lifespan 19.9 ± 0.5 days compared with 16.7 ± 0.4 days for controls) (Fig. 4K). In addition, they showed higher resistance to chronic ER stress (Fig. 4L). The increase in lifespan was also observed in another independently generated *manf-1* overexpression strain (*manf-1p::manf-1*, termed as *MANF-1^HAR^;* mean lifespan 18.5 ± 0.9 days compared with 15.9 ± 0.5 days for controls) (Supplementary Figure 5). Overall, these results demonstrate the multiple beneficial effects of MANF-1 in *C. elegans*.

### MANF-1 is secreted and localizes to lysosomes independently of the ER retention signal

Previously, we reported that *manf-1* is ubiquitously expressed in tissues such as the pharynx, hypodermis, and intestine [22, 23]. To further expand on these studies, we sought to provide novel insight into the mechanism of *manf-1* function. Moreover, we reasoned that altering the ER retention capability of MANF would be a suitable target for further investigation. For MANF to localize to ER, the protein must have a signal peptide at the N-terminus and a KDEL sequence (KEEL in worms) at the C-terminus. We generated several new transgenic strains expressing MANF-1::mCherry chimeric proteins using an endogenous promoter. Two such strains contained either an obstructed or deleted retention signal with mCherry fused at the C-terminus (*MANF-1^KEEL::mCherry^*and *MANF-1^ΔKEEL::mCherry^*, respectively) and one carried an unobstructed and functional ER retention signal after mCherry (*MANF-1^mCherry::KEEL^*). These different strains allowed us to investigate the subcellular localization of MANF-1 and how it might confer a protective response in animals.

The examination of *MANF-1^KEEL::mCherry^* and *MANF-1^ΔKEEL::mCherry^*animals with putatively secreted MANF protein revealed that the chimeric protein was present throughout the body, including the hypodermis, pharynx, coelomocytes, and the extracellular space (Fig. 5A–F’ and Supplementary Figure 6). A detailed examination identified puncta-like structures in the hypodermal cells as lysosomes based on colocalization studies. Specifically, the MANF-1::mCherry pattern overlapped with the lysosomal dye LysoTracker Green and the lysosomal membrane marker *scav-3::GFP* [29] (Fig. 5G). Similar to the hypodermis, MANF-1 localization in coelomocytes was confirmed to be lysosomal by colocalization with the marker *lmp-1::GFP* [30] (Fig. 5G). These data provide the first evidence that MANF-1 is secreted and localized to subcellular structures in *C. elegans*.

**Figure 5.**
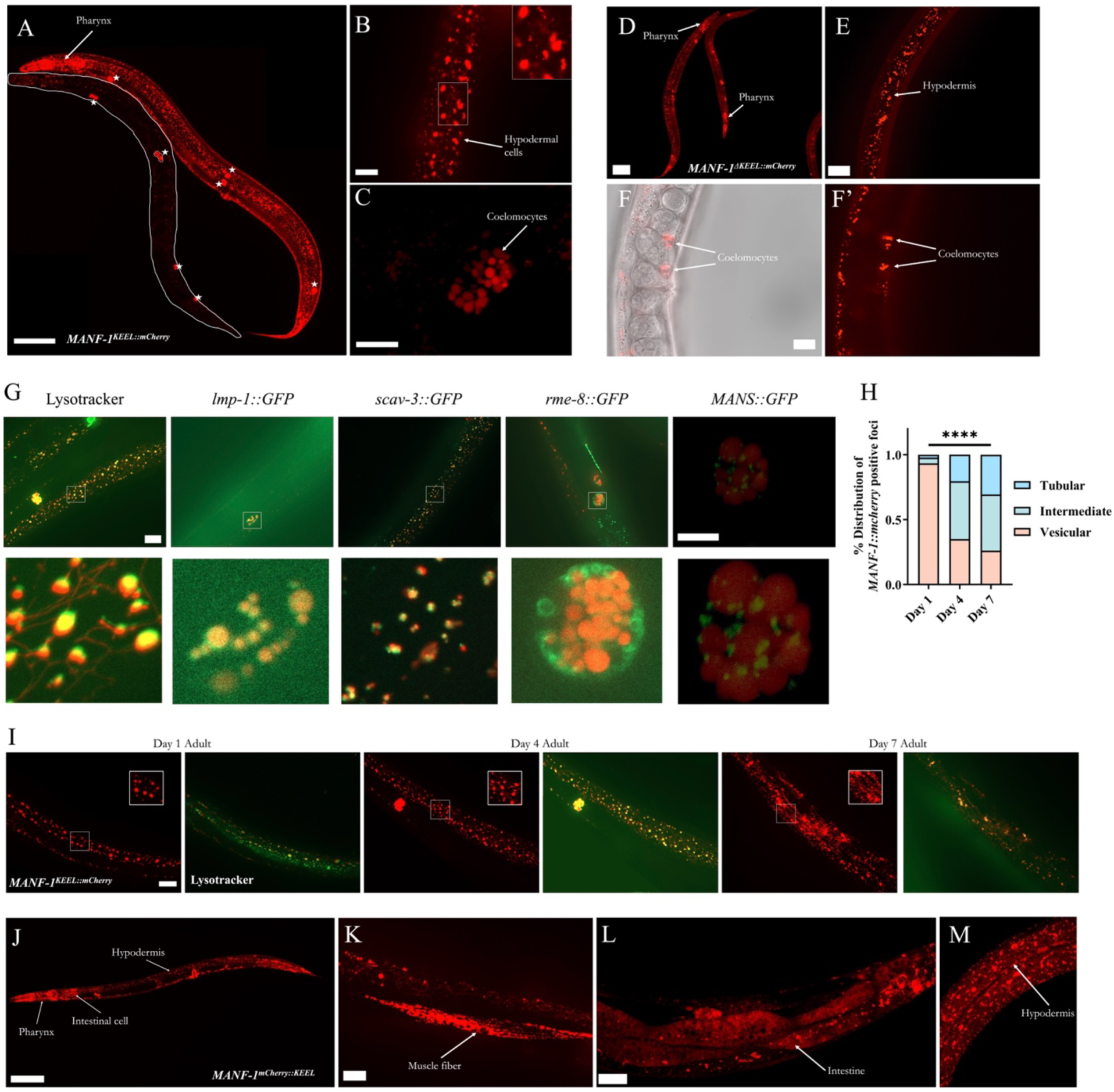
MANF-1 expression pattern and colocalization in *MANF-1^KEEL::mCherry^* transgenic animals. **A)** Two *MANF-1^KEEL::mCherry^*animals at day 1 of adulthood. The hypodermal region is in focus. MANF-1::mCherry fluorescence is throughout the body including the pharynx and coelomocytes (stars). The outlined worm has a weak fluorescence compared to the worm next to it. Scale bar 100 µm. **B-C)** Enlarged areas of *MANF-1^KEEL::mCherry^*animals showing fluorescence in hypodermal cells in clusters of varying sizes (scale bar 20 µm) (B) and in coelomocytes (scale bar 10 µm) (C). **D-F)** *MANF-1^ΔKEEL::mCherry^*animals at day 1 of adulthood. Fluorescence is similar to that seen in the *MANF-1^KEEL::mCherry^*line and observed in the pharynx (scale bar 100 µm) (D), hypodermal cells (scale bar 20 µm) (E), and coelomocytes (scale bar 10 µm) (F and F’). **G)** MANF-1::mCherry colocalization studies at day 1 of adulthood. Scale bars are 20 µm (top left-most panel, same for *lmp-1::GFP*, *scav-3::GFP*, and *rme-8::GFP*) and 10 µm (top right-most panel). The boxed region in each of the top row panels has been enlarged and shown below. MANF-1::mCherry chimeric protein was colocalized with LysoTracker™ Green DND-26 (hypodermal cells) and lysosomal membrane protein markers LMP-1::GFP (coelomocyte) and SCAV-3::GFP (hypodermal cells). No colocalization was observed with the endosome marker RME-8::GFP (coelomocyte) and golgi body marker MANS::GFP (coelomocyte). MANS::GFP animal was imaged using a Leica Confocal. **H)** Quantification of MANF-1::mCherry fluorescing structures. The foci were classified as vesicular, intermediate, or tubular and plotted as a stacked histogram. Worms were scored in at least three batches with 10 per batch. Data was analyzed using Chi-squared test. *p<0.05; **p<0.01; ***p<0.001; ****p<0.0001. **I)** *MANF-1^KEEL::mCherry^*animals at day 1, 4 and 7 of adulthood (scale bar 20 µm). Each animal has been imaged for MANF-1::mCherry alone (red) and MANF-1::mCherry with lysotracker (green) showing . In all cases hypodermal cells near the posterior region are shown. Insets show zoomed in view of MANF-1::mCherry foci that change from vesicular to tubular-looking structures. **J-M)** *MANF-1^mCherry::KEEL^*animals showing MANF-1::mCherry chimeric protein expression. Fluorescence in this animal is visible in the pharynx, hypodermis, and one of the intestinal cells (J) (scale bar 100 µm). (K) Zoomed in view of MANF-1::mCherry in a muscle fiber (scale bar 20 µm). Confocal images of an animal showing fluorescence in the intestinal (L) and hypodermal (M) regions. MANF-1::mCherry appears diffused with lysosomes appearing as bright fluorescing dots.

An identical expression pattern was observed in two additional independently generated *MANF-1^KEEL::mCherry^* strains with lower amounts of *manf-1::mCherry* (see Methods) (Supplementary Figure 7A, B). The MANF-1::mCherry positive puncta in these and the *MANF-1^ΔKEEL::mCherry^* strain were indistinguishable (Supplementary Figure 7C, D). Similar to *MANF-1^KEEL::mCherry^*, *MANF-1^ΔKEEL::mCherry^* animals showed significant lifespan extension and ER stress resistance (Supplementary Figure 7E, F). Another transgenic strain NK2548 that carries a *mKate2* reporter inserted at the 5′ end of the endogenous *manf-1* (downstream of the signal sequence, *manf-1p:: mKate2:: manf-1*), [23] also exhibited lysosomal localization of mKate2::MANF-1 upon treatment with tunicamycin (Supplementary Figure 8A). Additionally, detailed examination of both the mKate2::MANF-1 and the previously existing *manf-1p::GFP* transcriptional line also showed coelomocyte localization (Supplementary Figure 8A, B).

Further characterization of *manf-1* expression during aging revealed a dynamic pattern. As reported earlier for lysosomes [31], MANF-1::mCherry structures changed with age from a vesicular to tubular-like morphology ((Fig. 5H & I). No obvious overlap with RME-8::GFP (endosomes) [32], MANS::GFP (golgi body) [30], and GFP::LGG-1 (autophagosomes) was observed [33] (Figs. 5G and Supplementary Figure 9A). Additionally, we did not observe MANF-1::mCherry in other regions such as ER (using *vha-6p::GFP::C34B2.10*) [34] (Supplementary Figure 9B), mitochondria using (mitoGFP) [35] (Supplementary Figure 9C), and lipid droplets (using Bodipy 493/503) (Supplementary Figure 9D). Finally, MANF-1::mCherry was not detected in the neurons based on the DA neuronal marker *dat-1p::YFP* and pan-neuronal marker *unc-119p::GFP* [36, 37] (Supplementary Figure 9E, F).

We also examined MANF-1::mCherry localization in *MANF-1^mCherry::KEEL^* animals that contains a functional ER retention signal sequence (Figs. 5J–M). The expression patterns of these strains were like the observations of the two secreted MANF strains, *MANF-1^KEEL::mCherry^* and *MANF-1^ΔKEEL::mCherry^*. In addition, we observed bright fluorescence in the intestine in a pattern resembling ER-like morphology, which is consistent with the role of *manf-1* in ER-UPR maintenance, and in other tissues, such as the spermatheca and muscles (Fig. 5K-M). Fluorescence was also observed in several neuron-like cells in the ventral cord (data not shown). Intestinal and muscle expression was faint in early-stage larvae but became more prominent from the late larval stage and during adulthood. Both spermatheca were visible in adults and coincided with the egg-laying stage of the animals. Expression in lysosome-like structures became more prominent as the animals progressed from early larval to adult stages.

Taken together, these data demonstrate that MANF-1 is broadly expressed and its presence in various tissues is affected by the ER retention signal. The secreted MANF-1 localizes to lysosomes within hypodermal cells, where it may interact with other proteins to regulate lysosome function and proteostasis. These findings lead us to conclude that MANF-1 confers protective benefits on animals beyond its previously described role in the ER.

### MANF-1 promotes autophagy and lysosome function

The expression pattern of *manf-1* along with the phenotypic studies of mutant and transgenic animals prompted us to use a transcriptomic approach to understand the changes in gene expression associated with various cellular and molecular processes. To this end, we examined the differentially expressed (DE) genes in *manf-1(tm3603)* and MANF-1^HAR^ animals. A comparison of upregulated DE genes in MANF-1^HAR^ (5,943) with downregulated genes in *manf-1(tm3603)* (776) identified an overlapping set of 388 genes that were enriched in various GO, KEGG, and WormCat terms, including lysosome, lipid metabolism, and metabolism (Fig. 6A and B). An inverse of this analysis, that is, genes downregulated in *MANF-1^HAR^*and upregulated *in manf-1(tm3603)*, revealed 16 genes that were linked to terms such as ER function and ER-UPR maintenance (Fig. 6C and D). These data further support the role of *manf-1* in ER-UPR and lysosomal maintenance.

**Figure 6.**
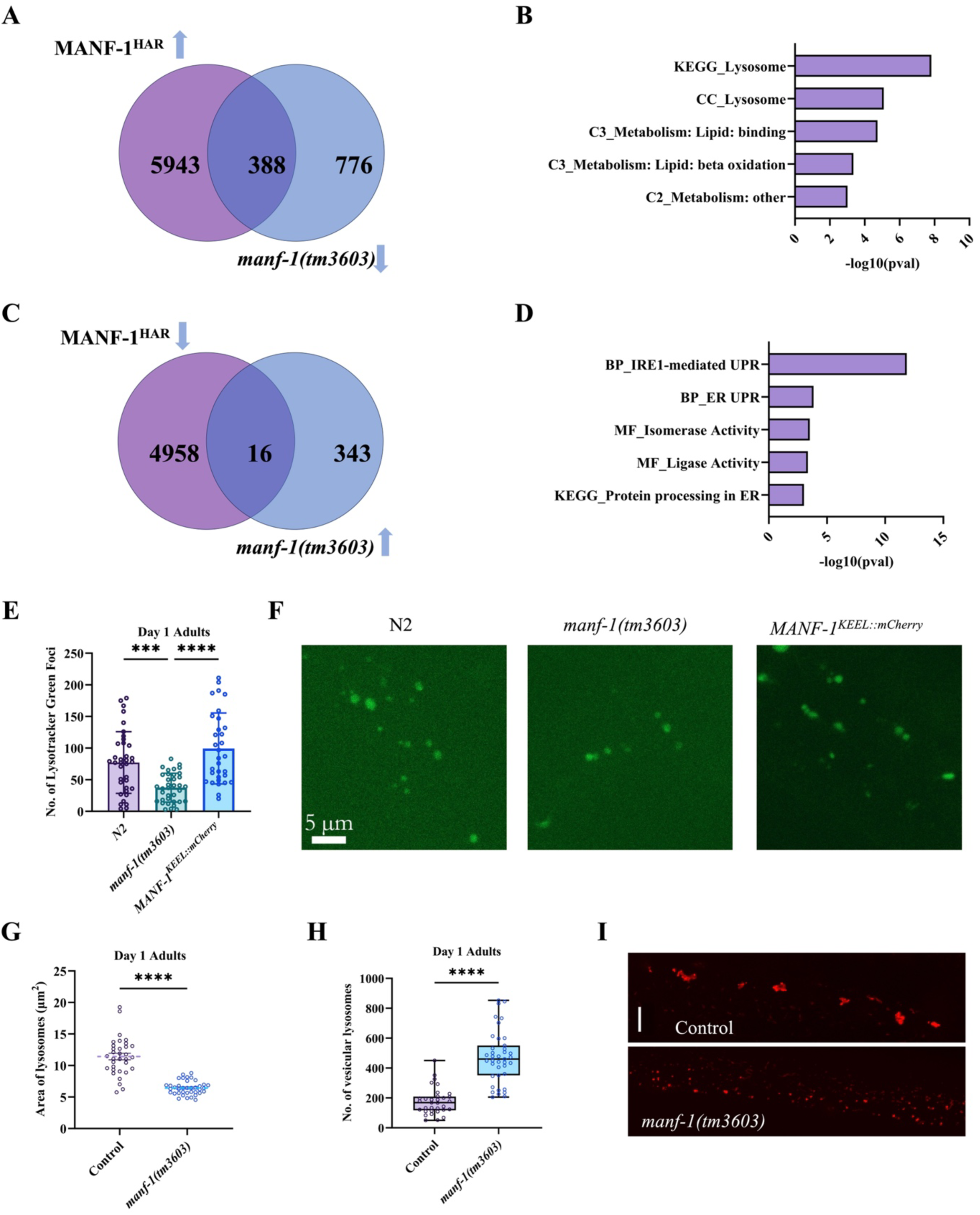
Transcriptome profiling of *manf-1* mutant and overexpression strains and the analysis of lysosomes. **A, B)** A total of 388 genes are common between the significantly upregulated genes in the MANF-1 overexpression line MANF-1^HAR^ when compared to the significantly downregulated genes in *manf-1(tm3603)* animals. Venn diagram (A) and Selected GO, KEGG and WormCat categories generated using easyGSEA from the eVITTA toolbox (B). **C, D)** A total of 16 genes are common between the significantly downregulated genes in MANF-1^HAR^ and significantly upregulated genes in *manf-1(tm3603)* animals. Data is plotted similar to panels A and B. **E-F)** Lysotracker staining of N2, *manf-1(tm3603)* and *MANF-1^KEEL::mCherry^* animals at day 1 of adulthood. (E) Quantification of total number of lysosomes stained. (F) Representative images showing Lysotracker-stained lysosomes. Scale bar 5 µm. **G-I)** Lysosomal size and count based on the *nuc-1::mcherry* reporter in *manf-1(tm3603)* and wild-type animals on day 1 of adulthood. Measurement of lysosomal area (G) and number (H) in the hypodermis of *nuc-1::mcherry* animals. (I) Representative Images corresponding to panels G and H showing changes in lysosomal size and area. Scale bar 10 µm. For panels E, G and H, at least three batches with 10-12 worms per batch were examined. Data is expressed as mean ± SD (E) and mean ± SEM (G, H). Panel H shows a box plot containing all data points along with the mean and lower and upper boundaries marking the 25^th^ and 75^th^ quartile, respectively. Data was analyzed using one-way ANOVA with Tukey’s test (E) and Student’s *t*-test (G & H). *p<0.05; **p<0.01; ***p<0.001; ****p<0.0001.

Prompted by the transcriptomic and expression data of MANF-1, we examined the lysosomes in *manf-1* null mutants. LysoTracker staining of *manf-1(tm3603)* animals revealed significantly fewer fluorescent structures when compared with N2 and *MANF-1^KEEL::mCherry^*adults (Fig. 6E and F). A lysosomal gene marker, *nuc-1::mcherry*, was also used [31], which revealed smaller lysosomes with a concomitant increase in their number in mutant worms (Fig. 6G–I). There may be multiple reasons for the differences in the LysoTracker and *nuc-1::mCherry* results. For example, *manf-1* mutant lysosomes may be less efficient regarding material uptake for degradation or less bright, making smaller lysosomes harder to see when stained with LysoTracker or less acidic, resulting in the dye being unable to stain adequately. The lysosomal defect in the *manf-1* mutant was further supported by increased tubular morphology in older adults (Supplementary Figure 10A).

We next examined the factors involved in the lysosomal and autophagic processes. Analysis of HLH-30/TFEB, a transcription factor that regulates autophagy and lysosomal genes in *C. elegans* [38], revealed enhanced nuclear localization in *MANF-1^KEEL::mCherry^* animals (Fig. 7A and B). As a positive control, we subjected animals to starvation, which resulted in a significant increase in HLH-30/TFEB being localized to the nucleus (Fig. 7A and B). Additionally, the expression of *hlh-30* was upregulated (Fig. 7C). Consistent with the essential role of *hlh-30* in mediating *manf-1* function, *hlh-30(tm1978); MANF-1^KEEL::mCherry^* animals had smaller MANF-1::mCherry puncta (Fig. 7D and E), a slower thrashing rate (Fig. 7F), and a shorter lifespan compared to the *MANF-1^KEEL::mCherry^* alone (Fig. 7G). These data indicate that *manf-1* utilizes *hlh-30* to exert cytoprotective effects in *C. elegans*.

**Figure 7.**
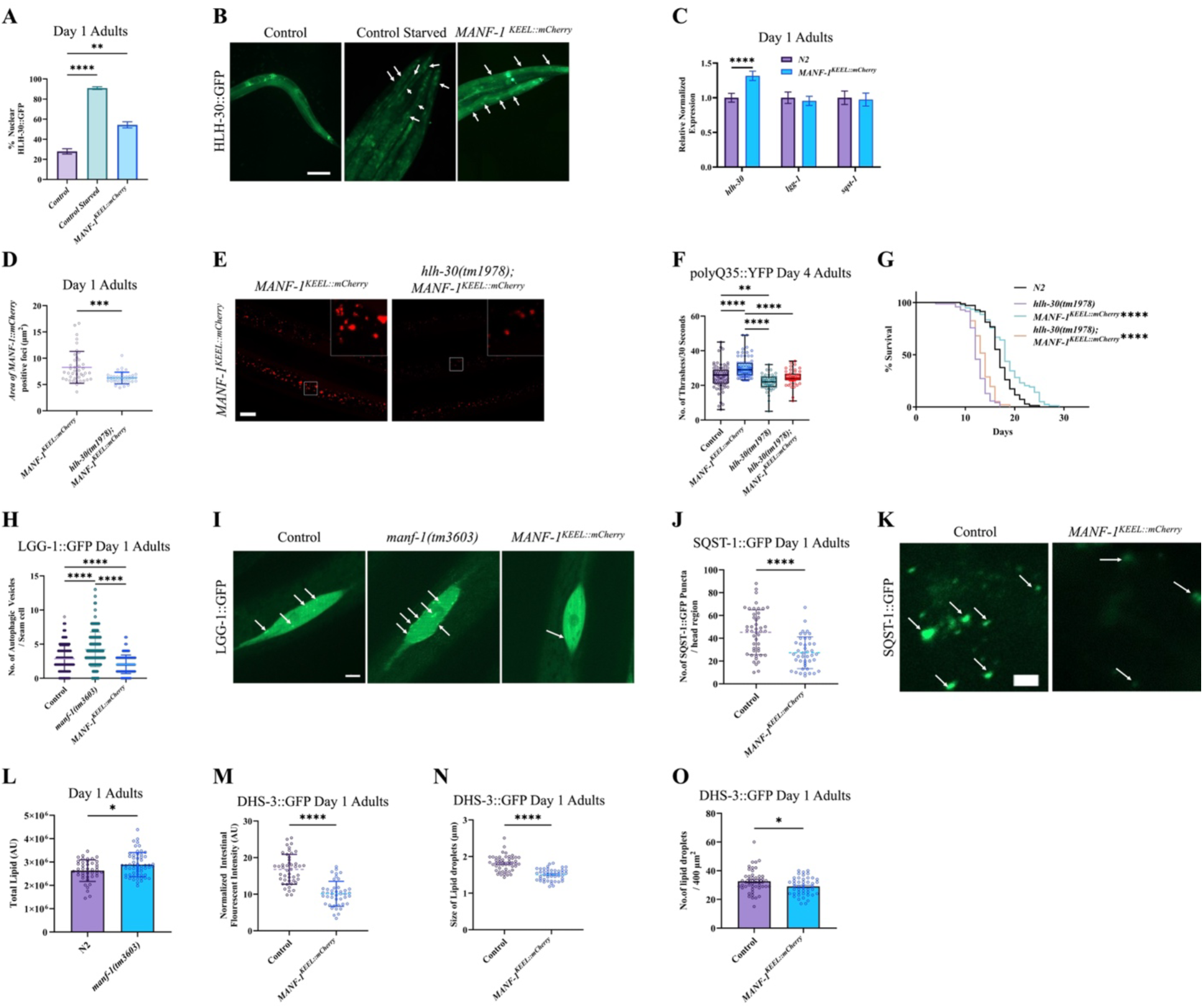
Analysis of HLH-30 expression, autophagy markers LGG-1 and SQST-1, and lipids in *MANF-1^KEEL::mCherry^* animals. **A)** Quantification of HLH-30::GFP nuclear localization in *MANF-1^KEEL::mCherry^* day 1 adults. Wild-type and starved animals were used as negative and positive controls, respectively. Graph is plotted as cumulative percentage of animals with nuclear localization. **B)** Representative fluorescent images corresponding to (A) showing HLH-30::GFP. Arrows point to areas of intestinal nuclear localization, scale bar 100 µm. **C)** RT-qPCR Analysis of *hlh-30, lgg-1* and *sqst-1* transcripts. **D)** MANF-1::mCherry foci size in the hypodermis. **E)** Representative images of MANF-1::mCherry foci corresponding to H, scale bar 20 µm. **F)** Thrashing rates of animals expressing polyQ35::YFP in a 30 sec interval. **G)** Lifespan analysis. **H)** Quantification of LGG-1::GFP puncta in seam cells. **I)** Representative images showing GFP fluorescence corresponding to panel H. Fluorescent puncta in LGG-1::GFP seam cells are visible (arrows). Scale bar 5 µm. **J)** Quantification of SQST-1::GFP puncta in the head region. **K)** Representative images corresponding to (J). SQST-1::GFP puncta in the posterior pharyngeal bulb. Arrows point to puncta in a 30 µm x30 µm region, scale bar 5 µm. **L)** Lipid quantification using Oil Red O staining. **M-O)** Lipid quantification in the intestine of using DHS-3::GFP. (M) Normalized GFP fluorescent intensity, (N) Lipid droplets and (O) number in a 400µM^2^ area. For the panel A, at least 2-3 batches with 30-40 worms per batch, panel C, three batches of pooled worms, and panels D, F-H, J, L-O, a minimum of three batches with 30-40 worms per batch were examined. Data are expressed as mean ± SEM (A, C, N, O) and mean ± SD (D, H, J, L, and M). Panel F shows a box plot containing all data points along with the mean and lower and upper boundaries marking the 25^th^ and 75^th^ quartile, respectively. Data was analyzed using one-way ANOVA with Dunnett’s test (A & H), Student’s t-test (C, D, J, L-O), one-way ANOVA with Tukey’s test (F), and log-rank (Kaplan-Meier) method (G). *p<0.05; **p<0.01; ***p<0.001; ****p<0.0001.

Two other autophagy genes, *lgg-1/LC3* and *sqst-1/*p62, were also examined. LGG-1 is involved in autophagosome formation and SQST-1 is an autophagy receptor that facilitates the degradation of ubiquitinated protein cargo [39, 40]. The results showed that although the *lgg-1/LC3* and *sqst-1/*p62 transcripts were unaffected in *MANF-1^KEEL::mCherry^* worms (Fig. 7C), a significant reduction was noted in LGG-1::GFP and SQST-1::GFP fluorescent puncta (Fig. 7H– K), consistent with increased autophagic clearance [41]. In agreement with this, *manf-1(tm3603)* mutants exhibited an increased number of autophagosomes (Fig. 7H and I). These findings led to the conclusion that the beneficial effects of MANF-1 are mediated by enhanced autophagy and lysosomal function.

As MANF-1^HAR^ animals showed enrichment of lipid metabolism genes and MANF-1 functions in an HLH-30-dependent manner, we investigated whether lipid content was also affected by HLH-30 based on its known role in lipid droplet clearance [42]. The results showed that *manf-1* mutants had elevated lipid levels (Fig. 7L and Supplementary Figure 11A), and the opposite phenotype was observed in *MANF-1^KEEL::mCherry^*animals, which included reduced amounts of lipids and smaller and fewer lipid droplets (Fig. 7M–O and Supplementary Figure 11B). Taken together, our data demonstrate that MANF-1 plays an essential role in the HLH-30-mediated pathway to promote autophagy and lysosomal function.

## DISCUSSION

The neurotrophic factor MANF was originally identified for its role in promoting dopaminergic neuron survival but subsequently was also found to affect other processes [4, 14, 15, 43, 44]. Much work on MANF has focused on its ER retention sequence, ER localization, and regulation of ER-UPR. Sequence and expression data suggest that MANF is an ER-resident protein released in response to ER stress and confers protection to neuronal cells. Although this aspect of MANF function has been studied in considerable detail, recent data suggest additional potential mechanisms that are not well understood. This is also supported by expression studies in animal models showing the presence of proteins in the cells of many tissues and extracellular regions [16, 21, 45–47].

We and others have previously reported that the *C. elegans* MANF homolog, MANF-1, plays an essential role in protecting dopaminergic neurons from increased ER stress [21–23]. In this study, we present a novel understanding of the function of MANF in the maintenance of the stress response, proteostasis, and longevity. We found that *manf-1* mutants exhibit chronic ER-UPR activation, and when subjected to ER stress, they exhibit enhanced neurodegeneration and a shorter lifespan. Additionally, both the transcript and protein levels of MANF-1 increased in response to ER stress, which is consistent with studies in other systems [9, 48–51]. As ER-UPR activation helps cells to manage stress through multiple mechanisms, including enhanced protein folding and clearance of nonfunctional proteins [7, 52], we examined whether *manf-1* mutant animals have defects in these processes. The results showed increased protein aggregation in PD and HD models that express human α-Synuclein and polyQ, respectively. These phenotypes were exacerbated in older adults and contributed to locomotor defects, suggesting an age-dependent breakdown of proteostasis in animals lacking *manf-1*.

Our work demonstrates that the overexpression of *manf-1* through different paradigms offers cytoprotective benefits and promotes healthy aging. We found that MANF-1 is expressed at different stages and in different cell types. Although there was no obvious overlap between MANF-1-expressing cells and DA neurons, the neurons may be protected by the uptake of the protein from the extracellular environment [21] since MANF-1::mCherry is seen at high levels in the pharynx and surrounding areas. Subcellular analysis of MANF-1::mCherry-expressing animals revealed that the protein is predominantly localized in the intestine, pharynx, hypodermal cells, and coelomocytes. The expression pattern observed within these transgenic lines showed tissue localization similar to the expression pattern of the endogenous *manf-1p::mKate2::manf-1* and the transcriptional *manf-1p::GFP* strains, where coelomocyte, intestinal, and pharyngeal expression was observed. Notably, we found that the expression pattern of MANF-1 was altered when the ER retention signal was blocked or deleted, resulting in a lack of localization in the intestine, muscles, and other areas, likely because the protein is secreted outside. This was consistent with previous studies involving cell cultures, which showed that removal of the ER retention sequence resulted in increased MANF secretion [45, 48]. The novel shift in MANF-1::mCherry expression in transgenic animals lacking ER signals, in conjunction with our results and rescue experiments, lead us to suggest that MANF-1 may be secreted from the intestinal ER and localizes to lysosomes of different cells to elicit its cytoprotective function. Furthermore, the identification of the intestine as likely synthesis point for MANF and the hypodermis as a target tissue for secreted MANF opens the door for future inquiry into the cell-non-autonomous function of MANF *in vivo*.

Our analysis of *C. elegans* MANF-1 was supported by transcriptomic data in which lysosomal genes were repressed in mutant animals and activated in those overexpressing *manf-1*. A previous study in *Drosophila* also identified lysosomal genes that were downregulated in the MANF mutant [47]. The authors also studied MANF localization in Schneider-2 cells, which revealed structures such as endosomes but not lysosomes. However, it is worth mentioning that they did not examine *in vivo* expression in whole animals. Overall, the transcriptomic data in flies and worms, along with our MANF-1::mCherry expression pattern, support the role of MANF in the lysosomal pathway to mediate its beneficial effect.

This study also provides the first evidence of MANF-1 expression in coelomocytes, specifically in their lysosomes. Proteins in coelomocytes, which are scavenger cells, play a role in the efficient uptake of materials from the pseudocoelom and endosomal to lysosomal trafficking [53]. Following uptake by coelomocytes, molecules travel through endocytic compartments and eventually enter lysosomes for degradation [54]. Although the exact mechanism underlying the role of MANF-1 in coelomocytes is unknown, it is likely to affect lysosomal function to maintain proteostasis. In support of this, Sousa et al. recently reported the GO enrichment of lysosomal and endosomal terms in MANF-deficient macrophages [55]. The authors also showed that recombinant MANF in older adults improved phagocytosis-induced lysosomal activity [55].

Recent studies have revealed that lysosomes act as signaling hubs inside cells, and the dysregulation of lysosomal function and autophagy are associated with various defects and neurodegenerative diseases [56, 57]. The autophagy-lysosomal pathway is instrumental in clearing toxic protein aggregates. We found that *MANF-1^KEEL::mCherry^* affects the gene expression and nuclear localization of HLH-30/TFEB, a transcriptional regulator of lysosomal and autophagy genes [58], which is necessary for lifespan extension conferred by Insulin/IGF-1 signaling and other longevity pathways [38]. HLH-30 is essential for the lifespan and proteostasis benefits of overexpressed MANF in MANF-1::mCherry animals. Previous studies have shown that the calcium released from lysosomes can activate calcineurin to dephosphorylate TFEB, leading to its nuclear localization [58]. Additionally, lysosomal calcium can trigger ER calcium release and signaling [56, 58]. These data, together with the observation that changes in ER calcium can promote MANF secretion [48], provide support to a model where MANF is part of a network that facilitates ER and lysosome crosstalk to regulate the TFEB-mediated activation of autophagy and proteostasis. Notably, we observed a reduction in the size of MANF-1-positive foci in *hlh-30* mutants, raising the possibility of a feedback mechanism. Consistently, the *manf-1* promoter contains sequences that overlap with the HLH-30 CLEAR-binding motif. Thus, similar to the lysosomal gene targets of TFEB [38], it is conceivable that *manf-1* is transcriptionally regulated by *hlh-30* and forms a positive regulatory network.

The role of MANF-1 in lysosome-mediated signaling and the involvement of HLH-30/TFEB in this process led us to investigate the role of LGG-1/Atg8/LC3 and SQST-1/p62 in regulating autophagy [40, 59–62]. Although an increase in LGG-1 and SQST-1 puncta may be caused by a blockage of autophagy, a reduction in both types of puncta is indicative of an increase in autophagic flux. We found that *MANF-1^KEEL::mCherry^*animals exhibited reduced LGG-1::GFP and SQST-1::GFP puncta, suggesting that MANF-1 promotes autophagy. Similarly, *manf-1(tm3603)* animals showed an increase in the number of LGG-1::GFP puncta. Consistent with these data, a recent study reported that MANF overexpression in mouse kidney cells promoted autophagy and mitochondrial biogenesis [41]. The authors also observed a decrease in p62 abundance and found that p-AMPK and FOXO3 play roles in promoting mitochondrial biogenesis. However, whether *C. elegans* AAK-2/AMPK and DAF-16/FOXO are involved in MANF-1-mediated processes remains unclear.

Our findings on HLH-30-mediated MANF-1 function were also supported by lipid analysis. As ER-UPR, autophagy, and lysosomes affect lipid metabolism [63–65], we assessed whether MANF-1 was also involved in this process. Examination of neutral lipids revealed increased levels in *manf-1* mutants. As expected, *MANF-1^KEEL::mCherry^* animals exhibited the opposite phenotype, which was accompanied by a reduction in the number and size of lipid droplets.

Earlier, TFEB was shown to be necessary for the clearance of lipid droplets and regulation of lipid catabolism [58]; in relation to this, mammalian TFEB and worm HLH-30 were found to regulate lysosomal genes and fusion of lipid droplets to lysosomes in a process called lipophagy [42, 66]. These data underscore the importance of MANF-1-HLH-30 signaling in multiple lysosome-mediated processes.

Our results have broad implications for understanding the function of MANF in higher eukaryotes and manipulating its role in promoting healthy aging and treat diseases including lysosomal disorders. In this regard, it is worth noting that *manf* expression is downregulated in Niemann-Pick and Gaucher disease models [67, 68]. Future work will help to understand the precise mechanisms of MANF localization, HLH-30/TFEB interaction, and stress response signaling that collectively promote neuronal health, proteostasis maintenance, and longevity.

## ACKNOWLEDGEMENTS

This work was supported by the Natural Sciences and Engineering Research Council of Canada Discovery grant to BPG and partially supported by the National Institutes of Health (R00-ES029552 to JHH). Some of the strains were obtained from the *Caenorhabditis* Genetics Center, which is funded by the NIH Office of Research Infrastructure Programs (P40 OD010440). The pGLC130 plasmid was made by Komal Prajapati. We thank WormBase for its citation database on individual genes (PMID 19910365) and the McMaster biology department confocal resource for assistance with imaging. Ram Mishra and Ravi Selvaganapathy provided feedback on some of the experiments. We thank Dr. Kacy Gordon (University of North Carolina-Chapel Hill) for her assistance with generating the MANF-1^HAR^ overexpression line.

## AUTHOR CONTRIBUTIONS

S.K.B.T. carried out the experiments and generated most reagents for the study. J.H. generated the *MANF-1^HAR^* strain and examined lifespan of animals. RNA-Seq experiments were carried out in J.H. laboratory. S.K.B.T., J.H. and B.P.G. analyzed the data. S.K.B.T wrote the first draft of the manuscript, S.K.B.T., J.H. and B.P.G. worked together to revise the manuscript. All authors reviewed the final version. B.P.G. supervised the study.

## COMPETING INTERESTS

The authors declare no competing interests.

## MATERIALS AND METHODS

### Strains and culturing

Strains were cultured on standard NGM (nematode growth media) agar plates using established protocols. Plates were seeded with *E. coli* bacterial strain OP50 as a food source [18]. Worms were grown and maintained at standard culture temperature (20°C) unless stated otherwise. Age-synchronized animals were obtained by using a standard bleaching protocol and treating gravid hermaphrodites with a mixture of sodium hypochlorite and sodium hydroxide (3:2::NaOCl:NaOH) [69]. The Bristol isolate of *C. elegans* (N2) was used as a wild-type control.

### Plasmid construction, site-directed mutagenesis, and transgenic strains

The plasmid pGLC72 (dat*-1p::YFP)* was described earlier [70]. Three others, pGLC130 (*hsp-16.41::manf-1*), pGLC180 (*manf-1p::MANF-1::mCherry*, carrying an obstructed or non-functional ER retention signal), and HAR002 (*manf-1p::manf-1*) were generated to study MANF-1 expression and its effect on cellular processes. For the pGLC130, a 507 bp cDNA fragment of *manf-1* was amplified by PCR using primers GL1041 and GL1042. The fragment was subcloned into pPD49.83 (a gift from Andrew Fire, Addgene plasmid # 1448; http://n2t.net/addgene:1448 ; RRID:Addgene_1448) using *BamH1* and *Nco1* restriction sites. The pGLC180 was created by PCR amplification of a 3,429 bp genomic fragment containing 2,713 bp of the *manf-1* promoter and *manf-1* gene (705 bp) using primers GL1749 and GL1750. The fragment was cloned into an *mCherry* reporter carrying vector, pHT101 (a gift from Casonya Johnson, Addgene plasmid # 61021; http://n2t.net/addgene:61021; RRID: Addgene_61021) using *Pst1* and *Age1* restriction sites. The third plasmid, HAR002, contains full-length *manf-1* genomic DNA driven by the *manf-1* promoter. The DNA fragment was subcloned into the Fire vector pPD95.75 using directional cloning upstream of the *unc-54* 3’UTR. The resulting plasmid was confirmed by sequencing.

Three additional *manf-1* plasmids were generated by site-directed mutagenesis that affect the ER retention signal sequence. This involved creating deletions and insertions in the gene using NEB Q5® Site directed mutagenesis kit (E0554). Mutagenic primers were designed using the online tool NEBaseChanger^TM^ and the provided annealing temperature was used for the PCR. As per the kit instructions, 1 ng/µL of pGLC180 was amplified using Q5 hot start high-fidelity 2x master mix and routine PCR thermocycling conditions. The PCR product was then incubated at 24°C for an hour in the NEB KLD enzyme mix and introduced into NEB DH5α cells by transformation. The plasmid pGLC189 (*manf-1p::MANF-1ΔKEEL::mCherry*) was created by deleting the 12 bp of native ER retention sequence (KEEL) of *MANF-1* in pGLC180 using primers GL1908 and GL1909. The pGLC189 was subsequently used as a base plasmid to generate pGLC196 (*manf-1p::MANF-1::mcherry::KEEL*) that carries 12 bp native ER retention sequence (KEEL). All plasmids were sequenced before generating transgenic strains.

The HAR002 plasmid was injected at 50 ng/µl concentration into the gonad of adult *unc-119(ed4)* animals along with the *unc-119* rescue plasmid (50 ng/µl), and 50 ng/µl EcoRI-cut salmon sperm DNA. Once extrachromosomal lines were established, plasmid was integrated by gamma irradiation as previously described [71]. The integrated line jhhIs001 was outcrossed with N2 to remove possible background mutations. All other plasmids were injected into the germline of adult N2 hermaphrodites using *dat-1p::YFP* (pGLC72; 25-50 ng/µL range) as a co-injection marker [70]. The pGLC130 (50 ng/µL) was used to generate two independent lines DY695 (*bhEx291*) and DY696 (*bhEx292*), both with genotype *hsp::manf-1; dat-1p::YFP*. One of these, DY695, was selected for detailed studies and RT-qPCR analysis showed roughly 15x increase in *manf-1* expression following 1 hr heat shock at 31°C. The pGLC180 (30 ng/µL) was used to generate three independent lines, DY757 (*bhEx302*), DY758 (*bhEx303*) and DY759 (*bhEx304*), with transcript levels ranging from 3x to 6x determined by RT-qPCR (data not shown). Additional transgenic animals were generated by injecting lower concentrations of pGLC180: DY798 (*bhEx305*) 10 ng/µL and DY800 (*bhEx306*) 1 ng/µL. The genotype of these animals was *manf-1p::manf-1::mCherry; dat-1p::YFP*. The pGLC189 was injected at 20 ng/µL to create DY807 (*bhEx307*), DY808 (*bhEx308*) and DY809 (*bhEx309*) strains with the genotype: *manf-1p::MANF-1ΔKEEL::mCherry; dat-1p::YFP.* The pGLC196 (25 ng/µL) plasmid was used to create DY819 (*bhEx310*) and DY820 (*bhEx311*) with the genotype: *manf-1p::MANF-1::mCherry:: KEEL; dat-1p::YFP*.

### RNA extraction and RT-qPCR

Total RNA was extracted from adults using Trizol (Sigma-Aldrich Canada, Catalog Number T9424). In brief, animals were bleach synchronized at desired adult stages. Worms were collected and washed with M9. Trizol was added followed by flash freezing in liquid nitrogen and then thawed in a 37°C water bath. The flash freeze and thaw steps were repeated three to four times. Chloroform was added, mixed, and then centrifuged to collect the aqueous phase. Isopropanol was used to precipitate RNA followed by two washes in 75% ethanol. Samples were allowed to air dry to remove any remaining ethanol. Extracted samples were treated with TURBO DNA-free™ Kit (Catalog Number: AM1907, ThermoFisher Scientific) according to manufacturer’s instructions. The resulting samples were used to obtain cDNA using the SensiFAST cDNA Synthesis Kit (Catalog Number BIO-65053, Meridian Bioscience) following kit instructions.

RT-qPCR was performed (in triplicate) using the Bio-Rad cycler CFX 96 and the SensiFAST SYBR Green Kit (Catalog Number BIO-98005, BIOLINE, USA). Gene expression levels were normalized to housekeeping genes *pmp-3*, *iscu-1* (*Y45F10D.4*) and *cdc-42.* qPCR involving extrachromosomal lines was done by enriching cultures for worms carrying the reporter of interest. For this, 30 fluorescent day 1 adults were picked and allowed to lay eggs overnight. The progenies were grown to gravid adults and bleached to obtain F2 worms. For the DY759 strain, 200 of these F2 fluorescent day 1 adult worms were individually picked and used for RNA extraction.

### RNAi

RNAi-mediated gene silencing was carried out using an established protocol in our lab [72]. Knockdowns were performed from the egg stage unless stated otherwise. Worms were fed with an empty plasmid (L4440)-containing *E. coli* HT115 culture or a gene-specific plasmid culture from the Ahringer library.

### Lifespan assay

All lifespan assays were carried out at 20°C on NGM agar plates seeded with *E. coli* OP50 bacteria as previously described [73]. Assays were repeated in triplicates. Each batch contained roughly 30 animals per strain. Worms were scored every day for survival starting from day 1 adulthood. They were transferred to fresh plates on alternate days until no progeny were seen. Cases involving escaping, bursting at vulva, and progeny hatching internally were censored. Survival curves were estimated using the Kaplan-Meir method (see the statistical analysis section).

### Heat shock treatments

Animals were grown at room temperature until young adulthood, they were then transferred to a 35°C incubator for 2 hrs and assayed for their survival 24 hrs later. The *hsp::manf-1* transgenic worms (DY695) were age synchronized and grown till adulthood. One-day old adults were heat shocked at 31°C for 1 hr. For lifespan assays, animals were treated every other day starting from day 1 until day 7 of adulthood.

### Normarski fluorescence microscopy and fluorescence quantification

Worms were mounted onto glass slides containing 2% agar pad. Animals were anaesthetized using 30 mM stock solution of NaN_3_, if necessary. Fluorescence was visualized using a Zeiss Observer Z1 microscope equipped with an Apotome 2 and X-Cite R 120LED fluorescence illuminator (unless stated otherwise). Images were also taken using a Leica SP5 confocal and are specified in figure legends. Zeiss Zen2 blue (https://www.zeiss.com/microscopy/en/products/software/zeiss-zen.html) and NIH ImageJ (http://rsbweb.nih.gov/ij/) software were used to capture and analyze images DA neurons were scored using a protocol that we described earlier [72] and refined in this study. Specifically, the *dat-1p::GFP* reporter was used to visualize neuronal cell bodies and dendrites [70]. To quantify neurodegeneration, we used the following four morphological categories: 1: normal-looking neuronal cell bodies and dendrites (wild-type pattern), 2: dendritic damage (blebs, puncta, faint appearance, missing, etc.) but cell bodies appearing normal, 3: one or more cell bodies missing, faint, or abnormally shaped but dendrites appearing normal, and 4: type 2 and 3 combined. At least 3 independent batches of animals were scored with a minimum of 20 animals per batch.

For *hsp-4p::GFP* strain, whole animal images were taken to quantify GFP fluorescence following different treatments. Using the polygon tool of ImageJ (http://rsbweb.nih.gov/ij/), the total fluorescence of a selected area was obtained. Samples were analyzed in triplicate, a minimum of n = 15 per batch.

To quantify α-Synuclein::YFP fluorescence in *unc-54p::α-Synuclein::YFP* animals, we used a protocol published earlier by our lab [22]. A fixed area of the head region with the pharynx in focus was selected and images were captured at an optimal exposure rate. Total fluorescent intensity was obtained using the polygon and measure tool of ImageJ. Samples were analyzed in triplicate, with a minimum of n = 15 per batch.

To quantify fluorescence intensity and no. of aggregates in *unc-54p::polyQ35::YFP* and *unc-54p::polyQ40::YFP* animals was analyzed similar to α-Synuclein::YFP as described above. Images were taken of the entire worm with the pharynx in focus. Total whole worm fluorescent intensity was obtained using ImageJ’s polygon and measure tool. To count aggregates, the “Find Maxima” function in ImageJ was used. Samples were examined in triplicate, with a minimum of n = 10 per batch.

The *hlh-30p::hlh-30::GFP* strain was used to count fluorescing intestinal nuclei in one-day old adults using published protocols [38]. As a positive control, animals were starved by transferring them to an unseeded NGM agar plate for 24 hrs that induced nuclear location of HLH-30. Samples were analyzed in triplicate, a minimum of n = 20 per batch.

The LGG-1::GFP puncta in *lgg-1p::LGG-1::GFP* animals were quantified using published protocols [39, 59, 61]. Z stack images at 63x magnification were taken at optimal slice intervals determined by the Zeiss Zen2 blue software. The maximum projection was acquired and the GFP puncta were counted in 4 seam cells per animal. Samples were analyzed in triplicate, a minimum of n = 10 per batch.

The SQST-1::GFP puncta in *sqst-1p::SQST-1::GFP* animals were examined using a protocol based on published studies [40, 59]. Z stack images were taken at optimal slice intervals determined by the Zeiss Zen2 blue software. A maximum projection of the Z stacks was obtained and GFP positive puncta were counted in the posterior pharynx of the head region using ImageJ “Find Maxima” function. Samples were analyzed in triplicate, a minimum of n = 10 per batch.

Lysosomes were examined in age synchronized animals at 63x magnification. Z stack images, ranging from 10 to 15 stacks per image, were taken at optimal slice intervals determined by the Zeiss Zen2 blue software. The lysosomal morphology in hypodermal cells was grouped into three categories: tubular, intermediate, and vesicular. A Chi-square test was used to analyze the data. For LysoTracker Green stained structures, the total number of fluorescent foci within the field of view were counted in the hypodermis of individual animals. In the case of MANF-1::mCherry and NUC-1::mCherry fluorescing structures, the maximum projection was used to count the number of foci and to quantify their morphology in the hypodermis. The number of foci within the field of view per animal were quantified using Cell Profiler [74].

The *dhs-3p::dhs-3::GFP* strain [75] was used to quantify lipid droplets in age synchronized one-day-old adults. Z-stack images were captured at 63x magnification. The images ranged from 8 to 12 stacks per image at optimal slice intervals determined by the Zeiss Zen2 blue software. As per the published protocol [76], a 400 µM^2^ region of interest was selected in the tail region of worms. All visible lipid droplets were counted, and their diameter was measured using the line tool in software. Samples were analyzed in triplicate, n = 14 animals per batch.

### LysoTracker™ Green staining

LysoTracker™ Green DND-26 dye (Catalog Number: L7526, ThermoFisher Scientific) was used to stain the lysosomes in age synchronized worms as described earlier [31]. A stock solution of 250 μM was made. Worms were washed with M9 and placed in a 1.5 mL microcentrifuge tube containing 80 μL of dye in M9 buffer (working concentration 60 µM). The tube was wrapped in an aluminum foil. Animals were placed in a rotator for 1 hr and then transferred to an aluminum foil wrapped NGM plate seeded with OP50 bacteria. The plates were placed in the 20°C incubator for 2 hrs and animals were imaged.

### Oil Red O staining

Neutral lipids in one-day-old synchronized adults were visualized by Oil Red O dye (Thermo Fisher Scientific, USA) staining using a published protocol [77]. Images were acquired using a Nikon 80i Eclipse Nomarski fluorescence microscope fitted with a Micropublisher 3.3 RTV color camera and Q-imaging software. ImageJ was used to quantify the images [77]. Samples were analyzed in triplicate, n = 15 to 20 animals per batch.

#### Bodipy 493/503 staining

*Bodipy 493/503* (Catalog Number: D3922, Invitrogen) was used to stain lipid droplets in age synchronized worms as described earlier [78]. Worms were washed in M9 buffer prior to a 15 min incubation in 4% paraformaldehyde fixative. They were then flash frozen in liquid nitrogen and thawed in a 37°C water bath. The freeze-thaw cycle was repeated three times. In the end, worms were washed three times with M9. A stock solution of 1mg/mL BODIPY 493/503in DMSO was diluted to a working concentration of 1 µg/mL. The animals were incubated in the BODIPY solution for 1 hr at room temperature, washed three times with M9, and imaged.

### Thrashing Assay

The thrashing rate of polyQ35::YFP animals was quantified using a published protocol [79]. Age synchronized day 4 adults were placed in 10 μL of M9 buffer and the number of thrashes in 30 seconds were counted. Samples were analyzed in triplicate, a minimum of n = 15 per batch.

### Chemical treatments

Age synchronized day 1 adults were used for chemical treatments unless stated otherwise. Animals were exposed to desired concentrations of drugs as described earlier [72]. For *manf-1* and *hsp-4* expression assays, animals were exposed to tunicamycin on culture plates. The duration and concentration are mentioned in figure legends. Acute treatments of tunicamycin (50 ng/µL) and paraquat (200 mM) were carried out in liquid by exposing animals for 4 hrs as described earlier [73].

Lifespan and *hsp-4p::GFP* reporter assays were performed using published protocols [25]. For lifespan, day 1 adults were transferred to plates containing 25 ng/µL concentration of the drug. Viability was scored daily. Samples were analyzed in duplicate, each batch containing a minimum of 30 animals. The *hsp-4p::GFP* animals were placed in a 1.5 mL microcentrifuge tube containing 25 ng/µL of tunicamycin in M9 buffer and incubated on a rotator for 4 hrs. Samples were analyzed in triplicate, each batch containing a minimum of 20 animals.

### RNA Sequencing Analysis

RNA-seq data was processed using the TrimGalore toolkit (http://www.bioinformatics.babraham.ac.uk/projects/trim_galore/) which employs Cutadapt [80] to trim low quality bases and Illumina sequencing adapters from the 3’ end of the reads. Only reads that were 20nt or longer after trimming were kept for further analysis. Reads were mapped to the WBcel235r95 version of the worm genome and transcriptome using the STAR RNA-seq alignment tool [81]. Reads were kept for subsequent analysis if they mapped to a single genomic location. Gene counts were compiled using the HTSeq tool [82]. Only genes that had at least 10 reads in any given library were used in subsequent analysis. Normalization and differential expression were carried out using the DESeq2 Bioconductor package [83, 84] with the R statistical programming environment. The false discovery rate was calculated to control for multiple hypothesis testing. Gene set enrichment analysis [85] was performed to identify gene ontology terms associated with altered gene expression for each of the comparisons performed. We used eVITTA to perform a gene enrichment analysis on the genes that were common between *manf-1 (tm3603)* and MANF-1^HAR^ [86]. Specifically, the easyGSEA tool generated the enrichment terms in Figures 6C and D.

### Statistical analyses

GraphPad Prism 9.5.1 was used to plot all the graphs and perform statistical analyses. The Student’s *t*-test or analysis of variance (ANOVA) with a multiple comparisons test were performed depending on the number of conditions and comparisons to be made. Data from repeat experiments were pooled and analyzed together. Graphs were plotted with either standard deviation (SD) or standard error of mean (SEM) as appropriate (see figures). RT-qPCR results were analyzed using CFX Manager 3.1 software (Bio-Rad, Canada; https://www.bio-rad.com/en-ca/sku/1845000-cfx-manager-software?ID=1845000), which performed an ANOVA or *t*-test. SigmaPlot 14 was used to calculate the mean lifespan using the log-rank (Kaplan-Meier) method.

## SUPPLEMENTARY DATA

**Supplementary Figure 1.**
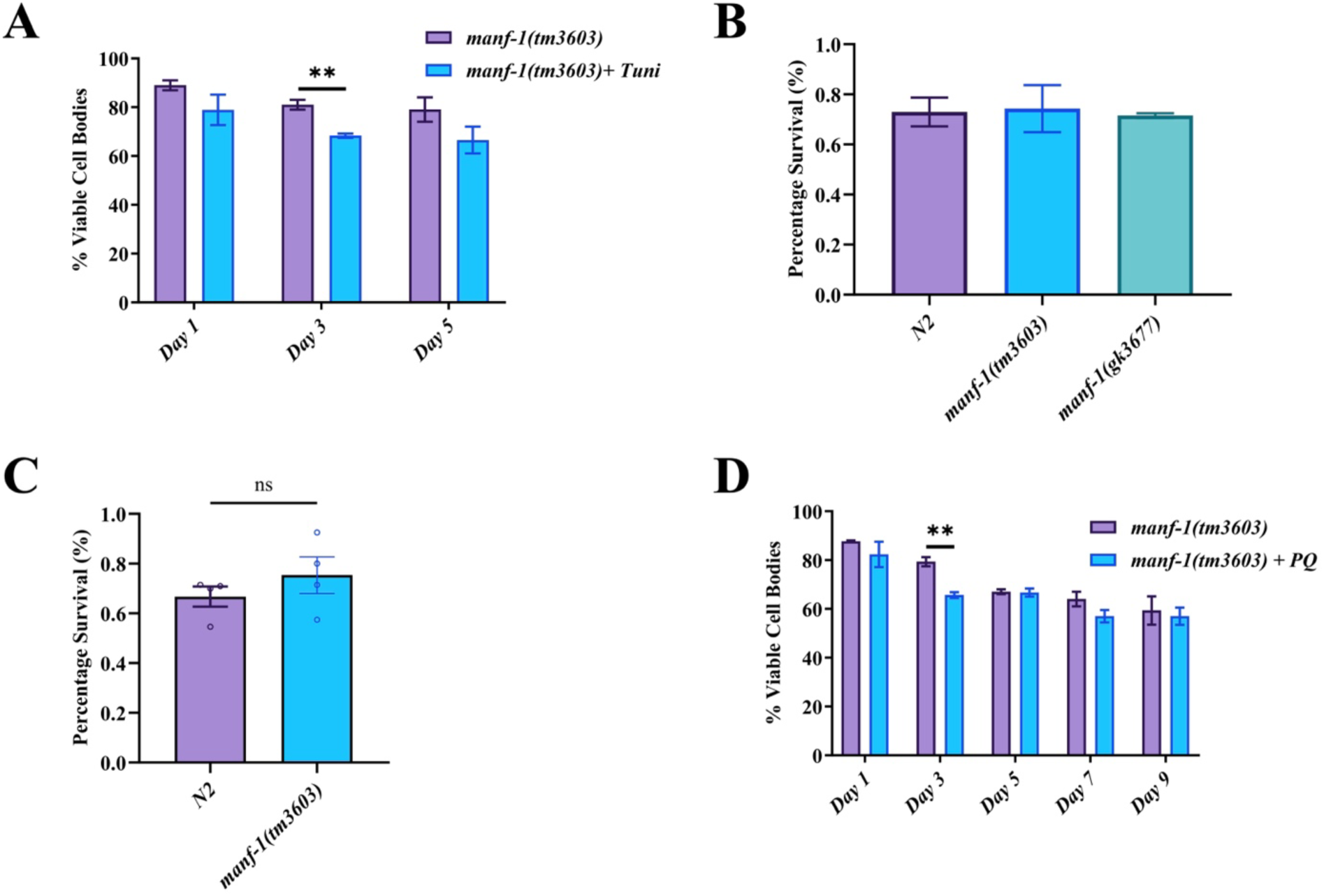
Effect of chemicals and heat on *manf-1* mutants. **A)** *manf-1* mutants were exposed to 5 μg/mL tunicamycin. The treatment reduced the percentage of viable DA neuron cell bodies by day 3. The neuronal damage plateaued by day 5. **B)** Survival of *manf-1* mutants following heat treatment at 35°C for 2hrs. Survival was examined after 24 hrs of recovery. *manf-1* mutants behave similarly to N2 worms. **C)** *manf-1* mutants were exposed to 200 mM paraquat for a 4 hrs period. **D)** *manf-1* mutants were exposed to 250 μM of paraquat. The treatment reduced the percentage of viable DA neuron cell bodies by day 3. For each assay, at least 3 batches of animals were examined with 10 - 30 per batch. Values are expressed as mean ± SEM. Data was analyzed using two-way ANOVA with Sidak’s test (A and D), one-way ANOVA with Tukey’s test (B), and Student’s *t*-test (C). *p<0.05; **p<0.01; ***p<0.001; ****p<0.0001.

**Supplementary Figure 2.**
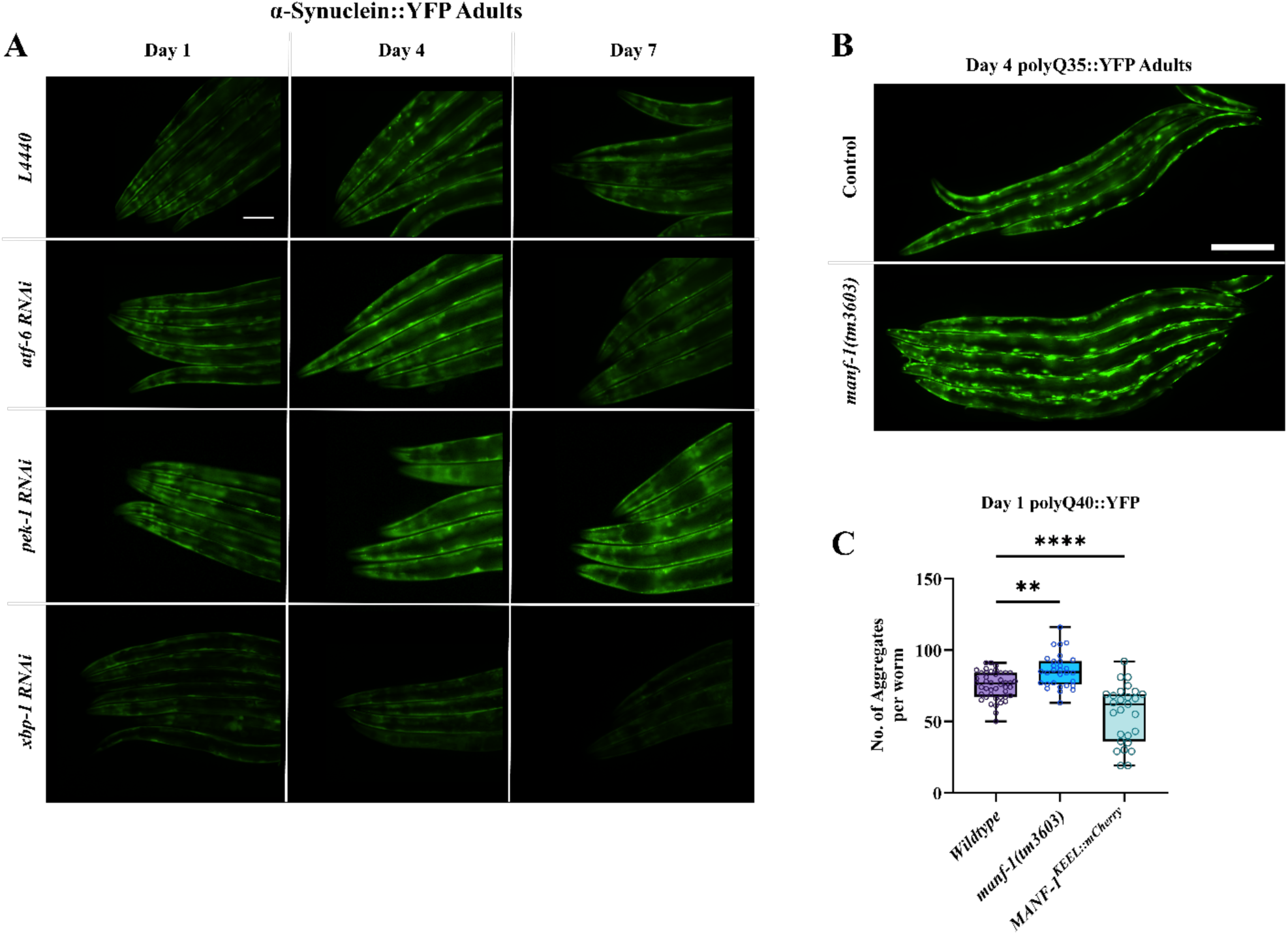
*manf-1(tm3603)* animals show increased aggregation of α-Synuclein::YFP and polyQ::YFP. **A)** Representative images of α-Synuclein::YFP fluorescence in *manf-1(tm3603)* animals measured on day 1, 4 and 7, following RNAi knockdown of *atf-6, pek-1* and *xbp-1*. L4440 refers to an empty vector control RNAi. Images correspond to panels D-F in Figure 2. Scale bar 100 µm. **B)** Representative images of polyQ35::YFP aggregation in *manf-1(tm3603)* and wild-type animals (control) on day 4 of adulthood. The images correspond to panels G-I in Figure 2. Scale bar 100 µm. **C)** Total number of polyQ40::YFP aggregates was significantly increased in *manf-1(tm3063)* but decreased in MANF-1^KEEL::mCherry^ day 1 adults when compared to wild-type controls. n = 15-20 worms per batch (2 batches). Box plots showing all data points along with mean and 25^th^ and 75^th^ quartiles. Data was analyzed using one-way ANOVA with Dunnett’s test or Student’s *t*-test. *p<0.05; **p<0.01; ***p<0.001; ****p<0.0001.

**Supplementary Figure 3.**
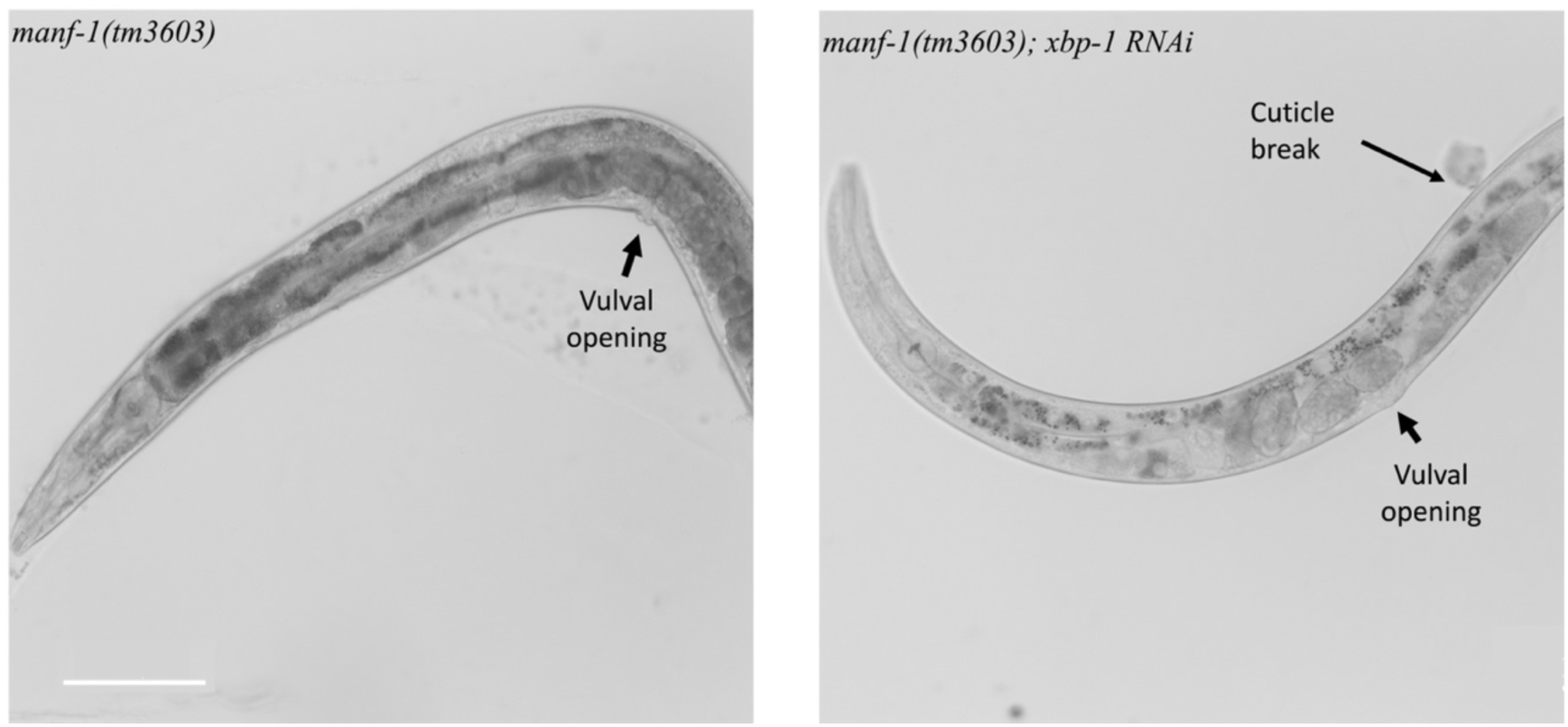
Effect of *xbp-1* RNAi knockdown on *manf-1* mutants. The *xbp-1* RNAi caused *manf-1(tm3603)* animals to become unhealthy. Representative images of day 1 adult animals are shown. The *manf-1(tm3603); xbp-1(RNAi)* animal is skinnier and more transparent compared to the *manf-1(tm3603)* control. In addition, the cuticle was weaker. The arrow points to an area where the cuticle was ruptured while mounting on the glass slide, causing internal contents to leak out. Head is towards the left in both panels. Scale bar is 100 µm.

**Supplementary Figure 4.**
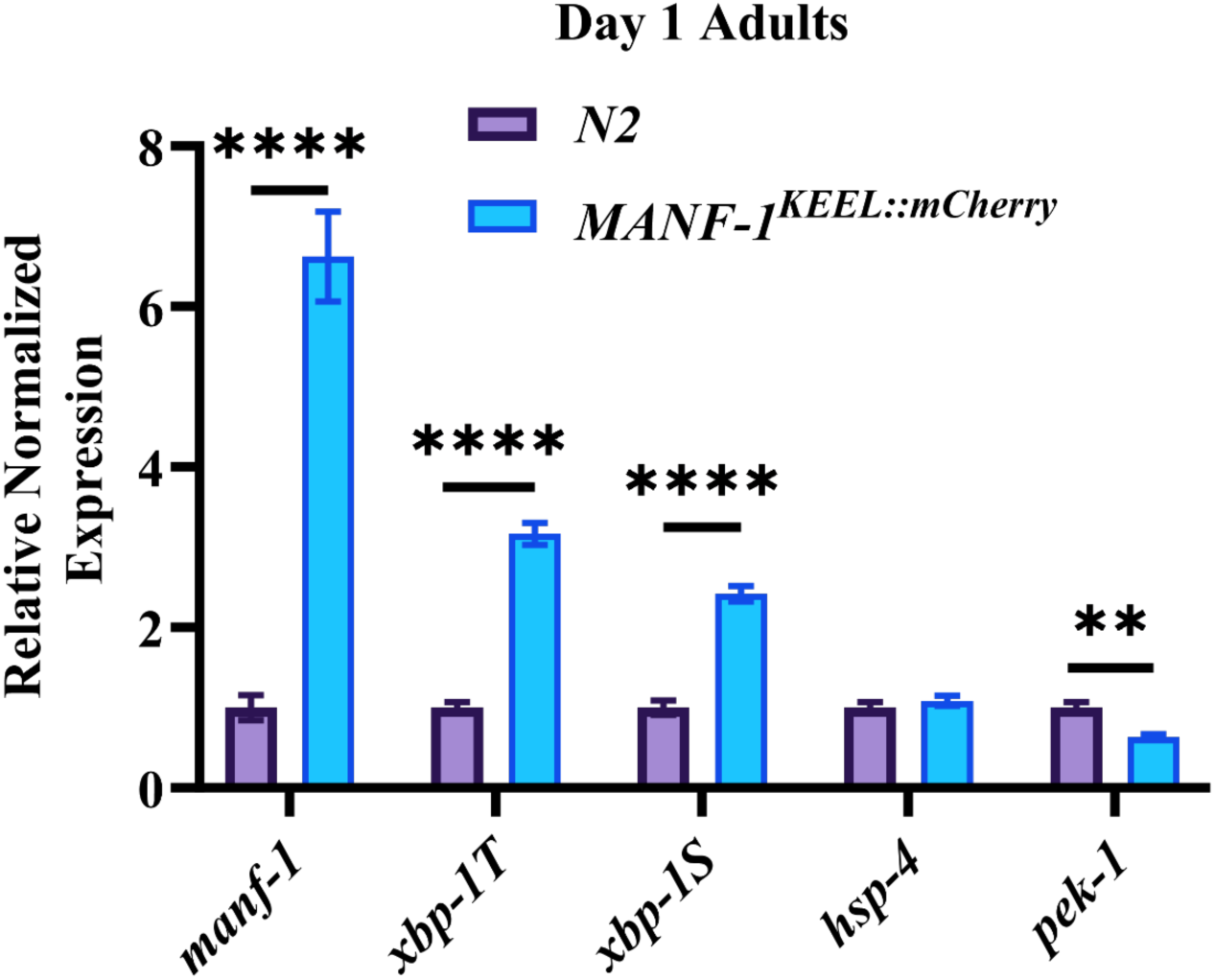
Transcript analysis of ER-UPR genes in *MANF-1* ^KEEL::mCherry^ animals. Expression levels were determined by RT-qPCR. **A)** *MANF-1* ^KEEL::mCherry^ had increased spliced *xbp-1 (xbp-1S)* and total *xbp-1 (xbp-1T),* but *hsp-4* gene expression was comparable to N2 and *pek-1* was reduced. The animals show ∼6.5 fold increase in *manf-1* transcripts. Results are expressed as mean ± SEM. Data was analyzed using student’s *t*-test. *p<0.05; **p<0.01; ***p<0.001; ****p<0.0001.

**Supplementary Figure 5.**
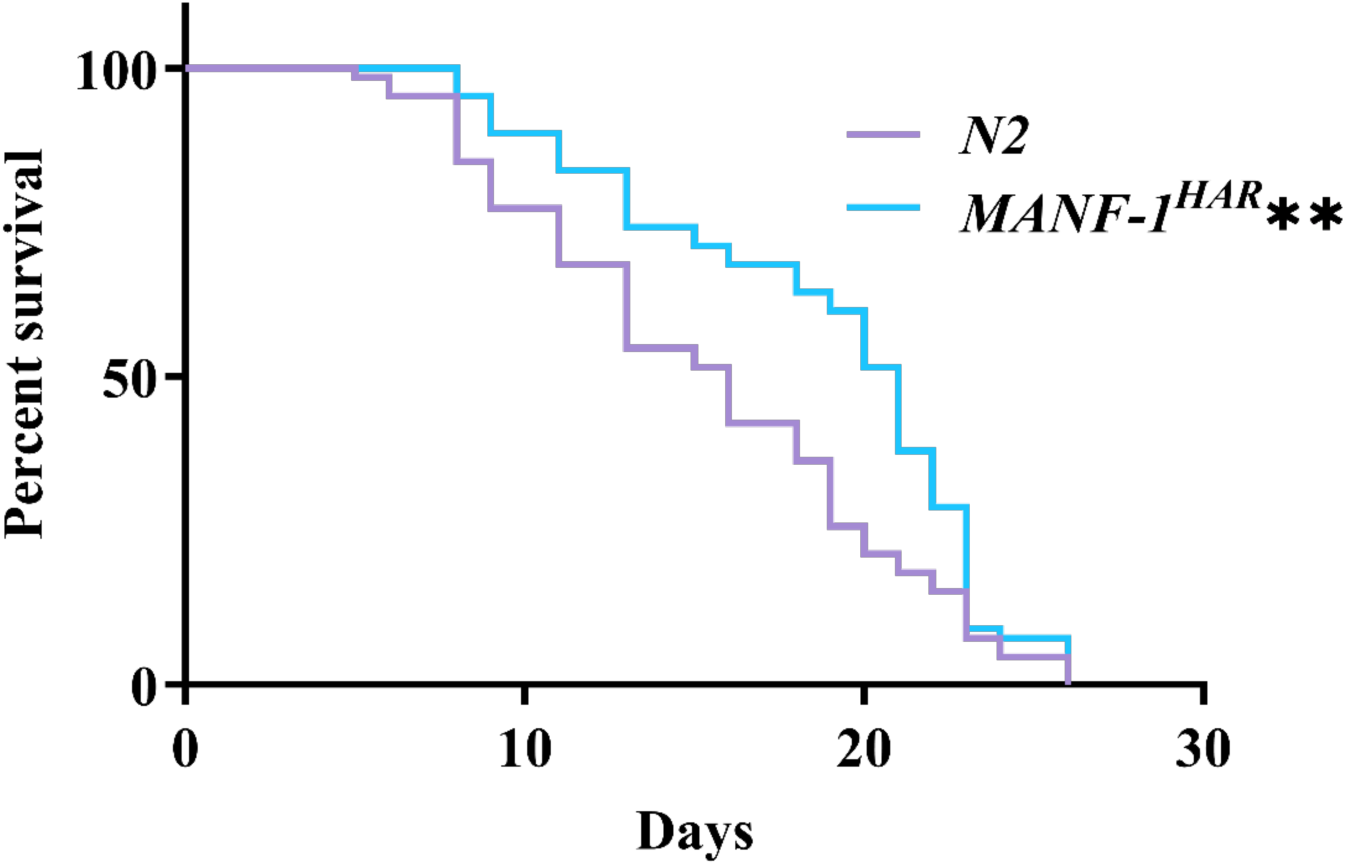
Lifespan analysis of MANF-1^HAR^ worms. The strain carries full-length *manf-1* gene under the control of the native promoter, which results in *manf-1* overexpression. The mean lifespan of MANF-1^HAR^ animals is 18.5 ± 0.9 days compared with 15.9 ± 0.5 days for controls. Approximately 30 worms were scored per batch and data from two independent batches was pooled. Statistical analysis was performed using the log-rank (Kaplan-Meier) method. *p<0.05; **p<0.01; ***p<0.001; ****p<0.0001.

**Supplementary Figure 6.**
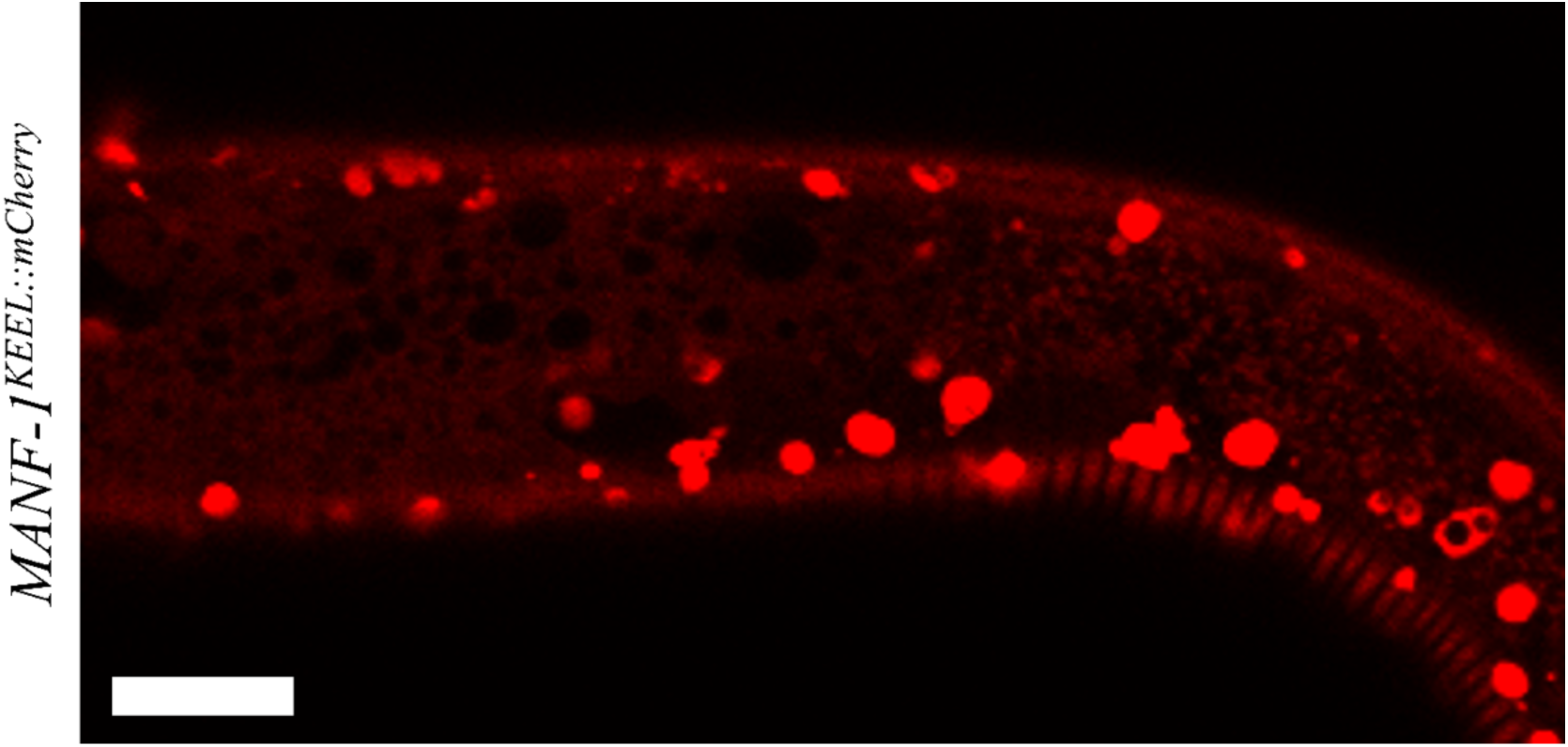
MANF-1::mCherry expression pattern. Confocal image of an *MANF-1^KEEL::mCherry^* animal showing lysosomes in hypodermal cells (bright structures) and diffused fluorescence throughout the body. Scale bar 10 µm.

**Supplementary Figure 7.**
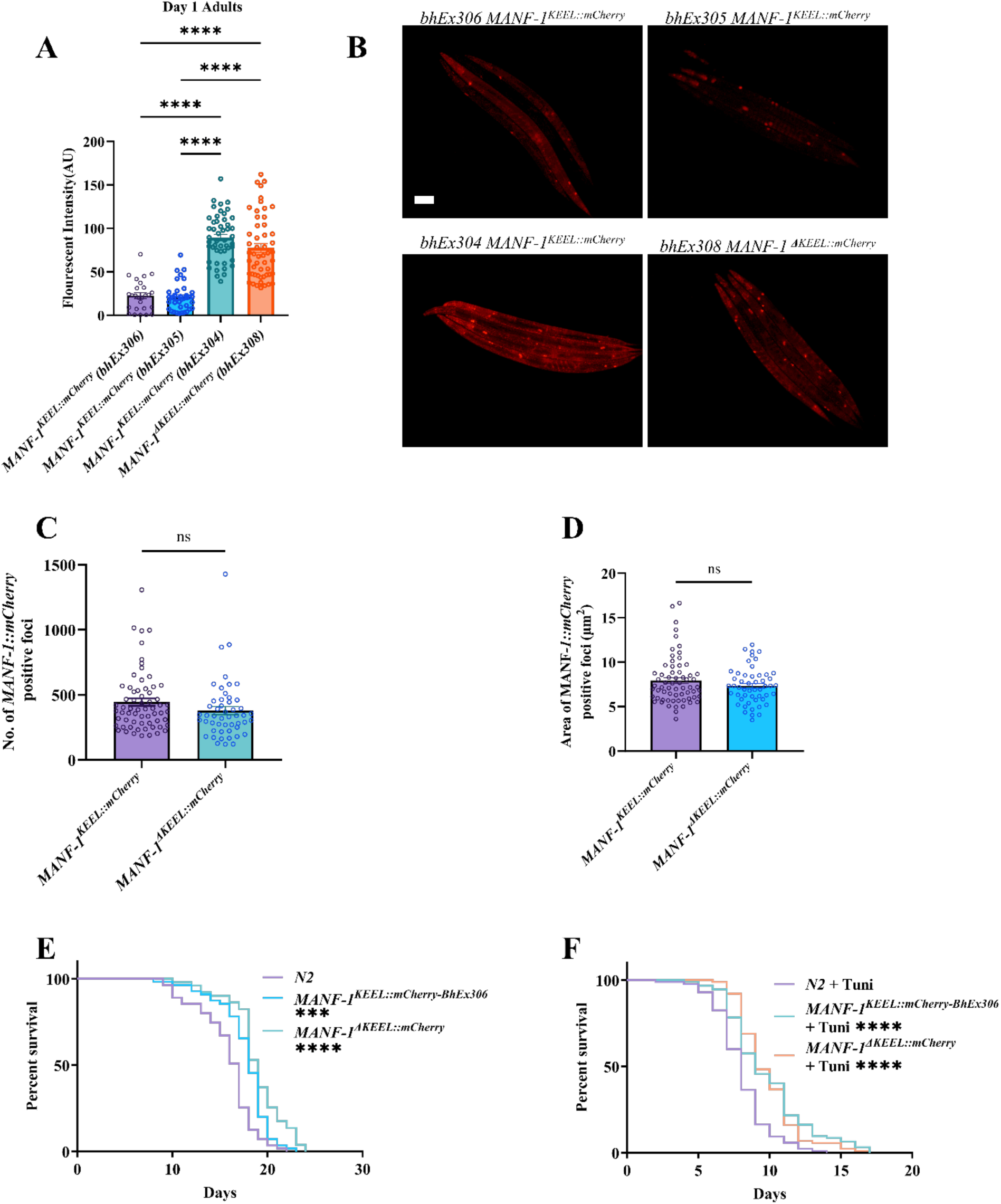
MANF-1::mCherry expression in various transgenic strains and lifespan phenotype of animals. **A)** Fluorescence intensity in *bhEx306* (pGLC180), *bhEx305* (pGLC180)*, bhEx304* (pGLC180), and *bhEx308*(*ΔKEEL*) transgenic worms at day 1 of adulthood. **B)** Representative images of animals corresponding to panel A. Scale bar 100 µm. **C, D)** MANF-1::mCherry foci number (C) and area (D) in day 1 adults of *MANF-1^KEEL::mCherry^ bhEx304* and *MANF-1^ΔKEEL::mCherry^ bhEx308* lines. **E)** Lifespan analysis of *MANF-1^KEEL::mCherry^ bhEx306 and MANF-1^ΔKEEL::mCherry^ bhEx308* at 20°C. Mean lifespans were as follows: N2 = 15.8 ± 0.408 days, *MANF-1^KEEL::mCherry^ bhEx306* 17.673 ± 0.376 days, and *MANF-1^ΔKEEL::mCherry^ bhEx308* 18.843 ± 0.414 days. **F)** Lifespan analysis of *MANF-1^KEEL::mCherry^ bhEx306 and* MANF-1^ΔKEEL::mCherry^ *bhEx308* following chronic tunicamycin exposure (25 ng/μL). Mean lifespan were as follows: N2 = 8.024 ± 0.212 days, *MANF-1^KEEL::mCherry^ bhEx306* 9.783 ± 0.299 days, *MANF-1^ΔKEEL::mCherry^*bhEx308 9.828 ± 0.227 days. A, C-F: n = 10 to 30 worms per batch (2 to 3 batches), Data in A, C, D is expressed as mean ± SEM. Comparisons were done using one-way ANOVA with Tukey’s test (A), Student’s t-test (C & D) and log-rank (Kaplan-Meier) method (E & F). *p<0.05; **p<0.01; ***p<0.001; ****p<0.0001.

**Supplementary Figure 8.**
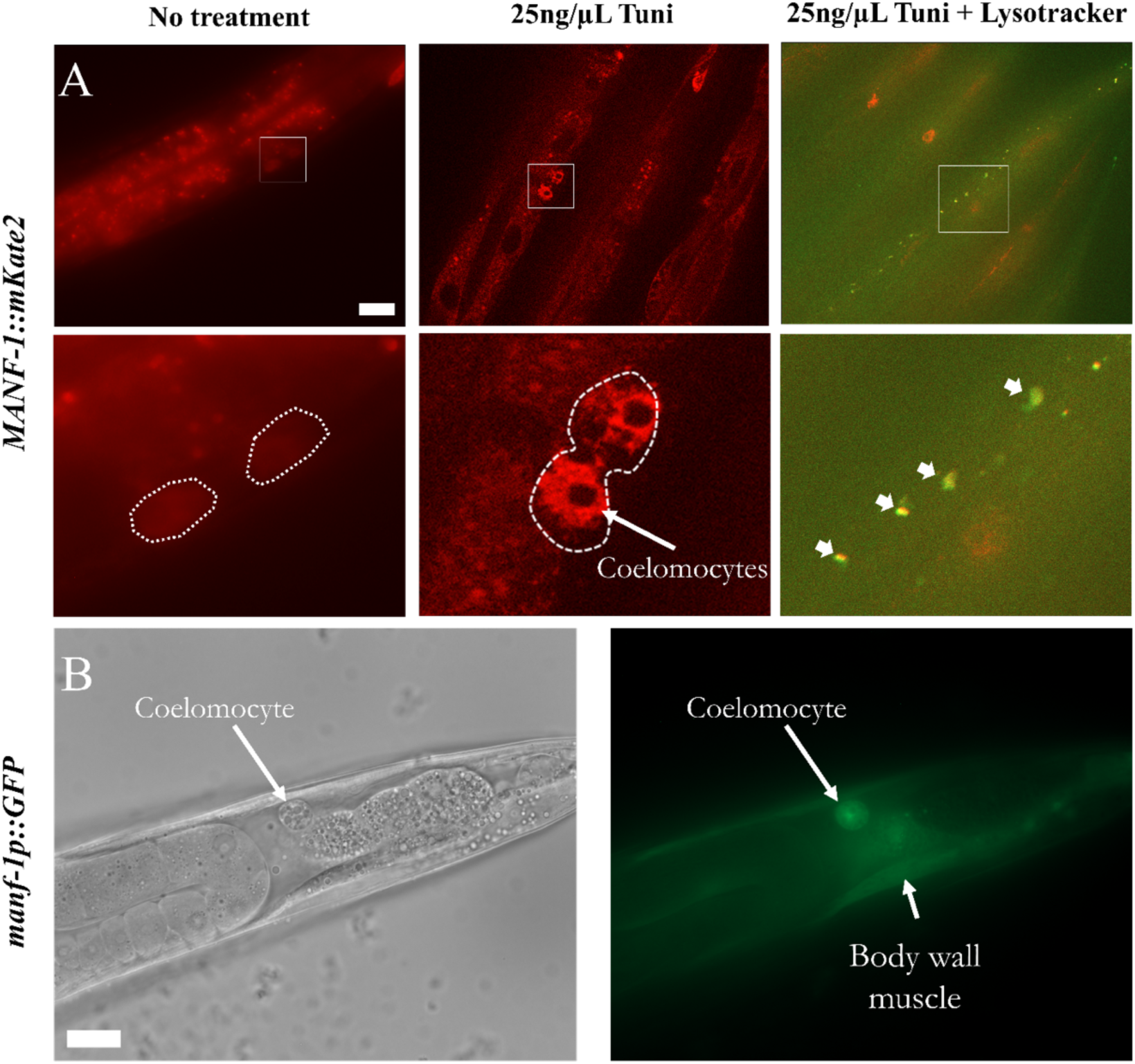
Reporter gene expression in *manf-1p::manf-1::mKate2* and *manf-1p::GFP* transgenic animals at day 1 of adulthood. **A**) The second row shows enlarged views of white boxed regions in the upper panel. The fluorescence is faint and diffused. Regions containing coelomocytes are marked. Treatment with 25ng/µL tunicamycin increased MANF-1 levels, thereby revealing an overlap with lysosomes (arrowheads). Scale bar 20 µm. **B)** Nomarski and GFP fluorescence images of a transcriptional *manf-1p::GFP* one-day old adult. The animal has GFP fluorescence in a coelomocyte and body wall muscle in the posterior region (arrows). Scale bar 20µm. Head is towards the left in all cases.

**Supplementary Figure 9.**
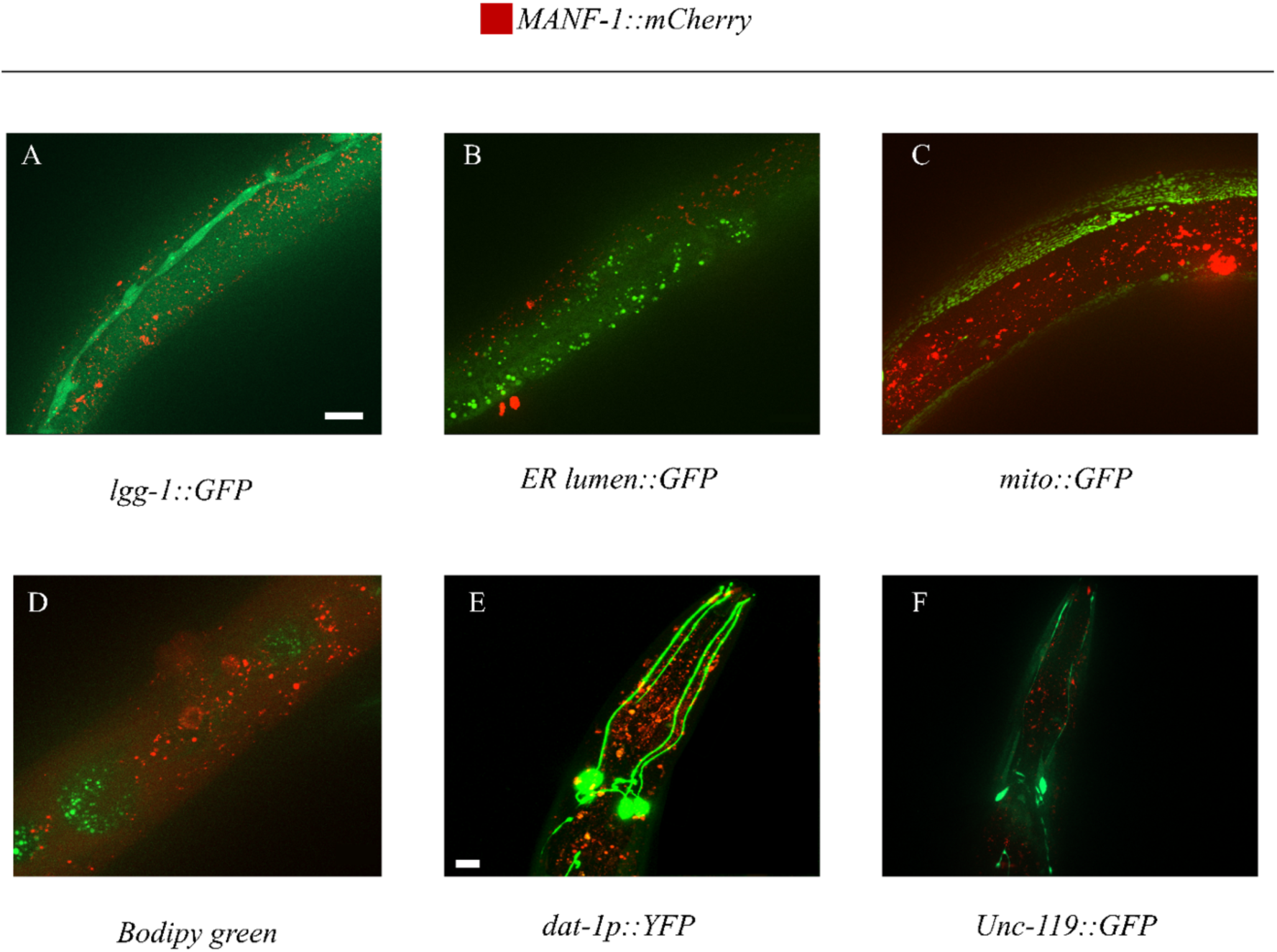
MANF-1::mCherry expression analysis in *MANF-1^KEEL::mCherry^* animals. Representative images are shown. **A-F)** No colocalization was observed with an autophagosome marker *lgg-1p::lgg-1::GFP* (A), intestinal ER lumen marker *vha-6p::GFP::C34B2.10(SP12)* (B), mitochondria marker *myo-3p::GFP*(mit) (C), lipid droplet marker *Bodipy green 493/503* (D), dopaminergic neuronal marker *dat-1p::YFP* ((E), and the pan-neuronal marker *unc-119p::GFP* (F). Scale bars in A-D and F are 20 µm. E was imaged with a Leica confocal microscope, scale bar 10 µm.

**Supplementary Figure 10.**
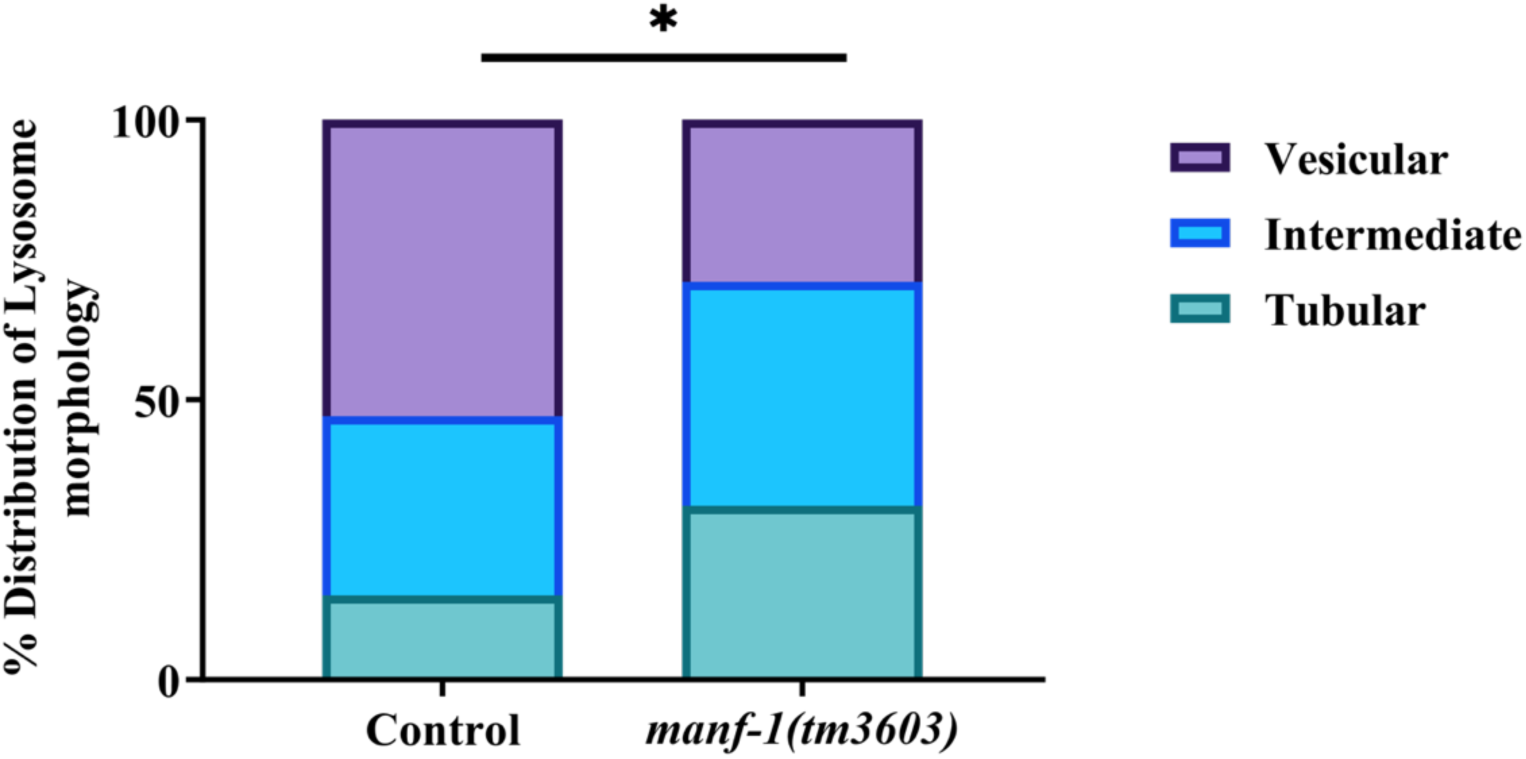
Loss of *manf-1* affects lysosomal morphology. Quantification of NUC-1::mCherry structures in *manf-1(tm3063)* four-day old adults. The lysosomes were analyzed and classified as vesicular, intermediate, or tubular and plotted as a stacked histogram. A total of three batches, n= 20 to 25 worms per batch, were plotted as stacked histograms. Data was analyzed using Chi-squared test. *p<0.05; **p<0.01; ***p<0.001; ****p<0.0001.

**Supplementary Figure 11.**
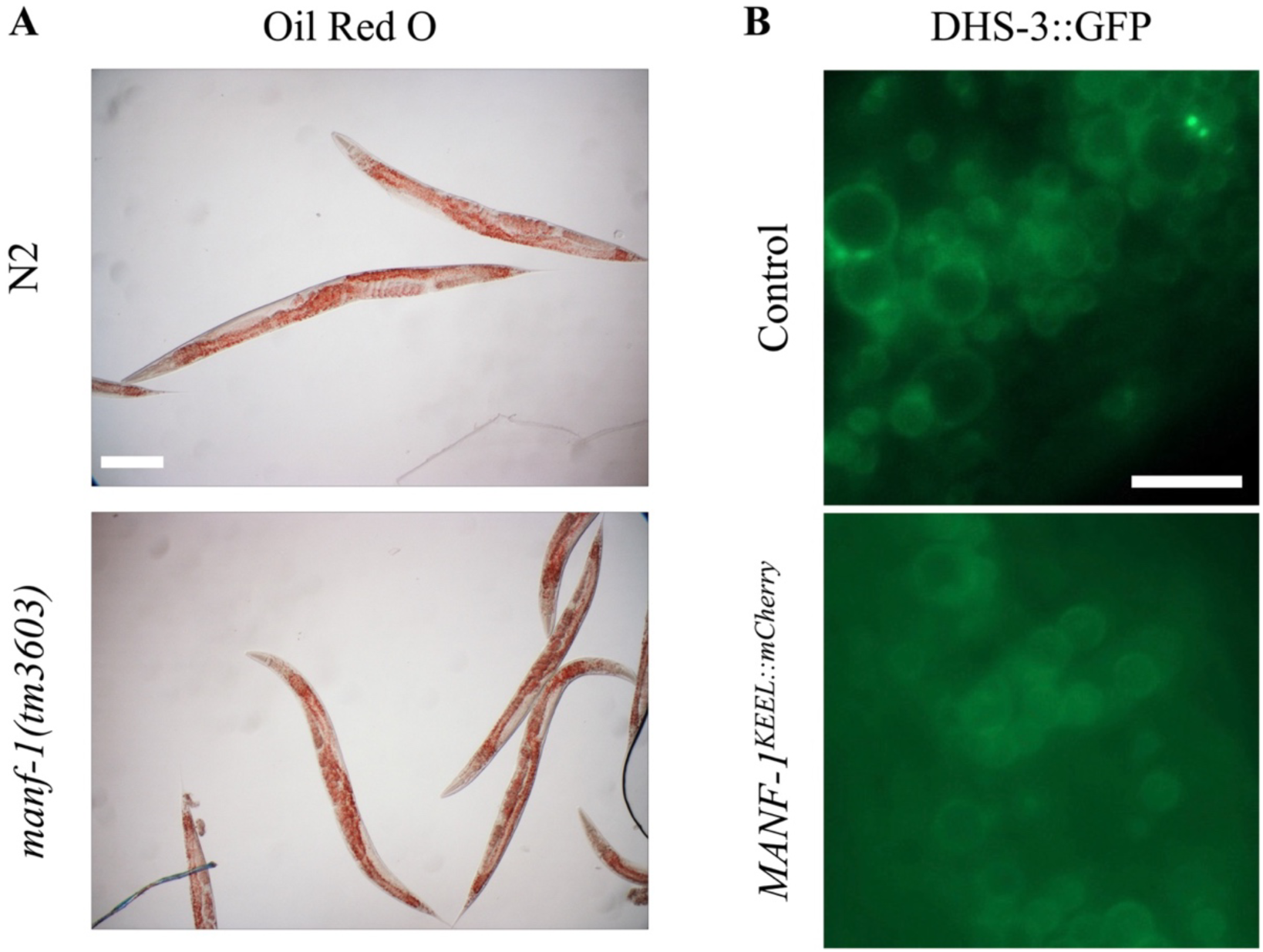
Representative images of animals showing Oil Red O staining and expression of the lipid droplet marker *dhs-3p::dhs-3::GFP*. (**A**) Oil Red O staining of day 1 *manf-1(tm3603)* mutants and N2. scale bar 10 µm. (**B**) Representative images of the lipid droplets in the tail region of day 1 *MANF-1^KEEL::mCherry^* adults, scale bar 5 µm.

## Notes

### Competing Interest Statement

The authors have declared no competing interest.

### Summary of Updates

The current version includes new results on MANF expression, mutant characterization, and transcriptomic study. Another author was added.

